# Mother Brain is Wired for Social Moments

**DOI:** 10.1101/2020.05.31.125955

**Authors:** Ortal Shimon-Raz, Roy Salomon, Miki Bloch, Gabi Aisenberg Romano, Talma Hendler, Yaara Yeshurun, Adi Ulmer-Yaniv, Orna Zagoory-Sharon, Ruth Feldman

## Abstract

Reorganization of the maternal brain, primed by oxytocin surge during childbirth, triggers the species-typical maternal social behavior. These brief social moments carry profound effects on the infant’s social brain and likely have a distinct signature in the maternal brain. Utilizing a double-blind, oxytocin/placebo administration crossover design, we imaged mothers twice while observing three naturalistic maternal-infant contexts in the home ecology; “unavailable”, “unresponsive”, and “social”, when mothers engaged in synchronous pick-a-boo play. We found four processes by which mother’s brain registers social moments. *Salience* - social moments increased activations throughout the maternal brain network; *Brain-behavior coupling* - caregiving behavior linked with socially-driven neural response; *Oxytocin sensitivity* – administration impacted neural response mainly to the social context; and *Temporal engrams*–consistent temporal patterns in insula and TP characterized response to social play. Findings describe how mother’s brain compiles and amplifies these precious social moments to generate dyad-specific brain-behavior patterns that initiate the cross-generational transmission of human sociality.

## Introduction

The maternal brain, the interconnected structures that reorganize upon the birth of an infant to enable mammalian mothers to care for infants, defines the key neural network sustaining mammalian sociality. Primed by the oxytocin surge during labor, the oxytocin-producing hypothalamus sends projections to sensitize a subcortical network that triggers expression of the mother’s species-typical social behaviors which usher young to social living (Feldman, 2015a; Numan and Young, 2016). These special moments of social contact reorganize the infant’s lifetime oxytocin system (Champagne et al., 2001; Feldman, 2016; Francis et al., 1999; Krol et al., 2019), augment the salience of social cues (Marlin et al., 2015), and sculpt the infant’s brain and behavior to life within the social ecology (Hammock, 2015). While few functions are as conserved as maternal care across mammalian evolution, in humans the subcortical structures comprising the maternal brain also include multiple insulo-cingulate, temporal, and frontal regions that coalesce to form the “human caregiving network” (Feldman, 2017). Activation of this network enables parents to perform the complex, multidimensional task of parenting human children; empathize with the infant’s emotion, mentalize to infer infant intentions, prioritize caregiving activities, and plan for long-term parenting goals based on culturally-transmitted social values (Feldman, 2015a, 2017).

Similar to other mammals, the human parental brain shapes and is shaped in turn by maternal behavior, particularly by moments of mother-infant synchrony - the temporally-coordinated rhythmic episodes of shared gaze, mutual positive affect, and “motherese” high-pitched vocalizations observed universally (Feldman, 2007). Albeit brief, these precious social moments expose mothers and infants to massive amounts of social inputs, require a temporally-fitted pattern to regulate the high arousal, and carry profound effects on infant development (Tronick, 1989). Longitudinal studies indicate that mother-infant synchrony builds dyad-specific patterns that maintain temporal consistency from infancy to adolescence and young adulthood (Feldman, 2010; Ulmer-Yaniv et al., 2020) and predict improved social-emotional outcomes (Feldman, 2020; Halevi et al., 2017). Moreover, activation of the parent’s “caregiving network” in the first months of parenting shapes child socioemotional competencies as mediated by parent-infant synchrony and parental oxytocin (OT) levels (Abraham et al., 2018, 2016; Kim et al., 2015). These findings highlight the links between the three components of human bonding: the parental brain, the OT system, and parent-child synchrony. Indeed, when maternal-infant bonding is disrupted, due to depression or environmental stress, deficits are observed in the parental brain, synchronous caregiving, and OT production, which carry long-term effects on child propensity to psychopathology and maturation of the social brain (Davis et al., 2017; Kim et al., 2016; Levy et al., 2019; Pratt et al., 2019).

Across species, the “maternal care” envelope marks the overall provisions transmitted from one generation to the next that contain the evolutionary acquired information necessary for survival and program the infant’s brain to what it means to be a member of that species (Kundakovic and Champagne, 2015; Meaney, 2001). Maternal care comprises a range of long and arduous activities, such as nest building, food retrieval, and, in some primate species, group collaborative and defensive activities (Hayes, 2000; Russell et al., 2003). Episodes of maternal social contact interfacing with an individual infant are brief, and, in some species, last no longer than several minutes per day for several consecutive days (González-Mariscal, 2007; Lucion and Bortolini, 2014). In humans, moments of direct mother-infant social contact are similarly brief and occupy a fraction of the overall maternal caregiving. Between three and nine months, episodes of mother-infant face-to-face synchrony typically last three to five minutes; however, their impact is pervasive and long-lasting (Cohn and Tronick, 1988; Feldman, 2015b). One mechanism sustaining the effects of these brief social moments is *bio-behavioral synchrony* (Feldman, 2017). Moments of mother-infant behavioral synchrony provide a template for the coordination of physiological processes, allowing the mature brain to externally-regulate the immature brain and tune it to social living (Hofer, 1994; Leong et al., 2017). During synchronous moments, mothers and infants coordinate their heart rhythms (Feldman et al., 2011b; Gray et al., 2017), oxytocin response (Feldman et al., 2010), and neural oscillations (Leong et al., 2017), and these episodes carry an “imprinting-like” effect on the infant’s social brain. It is thus likely that these intense social moments also have a distinct signature in the maternal brain.

What may be the mechanism by which the human maternal brain registers these precious social moments? We suggest four processes marking the special role of distinctly-social exchanges in the maternal brain; (1) salience, (2) brain-behavior coupling, (3) oxytocin sensitivity, and (4) “temporal engrams”. *Salience* describes the increased response of structures in the “human caregiving network” in response to prototypical social moments, including the subcortical structures (amygdala, VTA), and the human-specific para-limbic (AI, ACC), temporal (STS/STG, TP), and frontal (mPFC) structures (Feldman, 2015a; Kim et al., 2016; Swain et al., 2014). While the brain of any adult exhibits responses in this network to infant cues (Kringelbach et al., 2008; Rilling and Mascaro, 2017), the mother’s neural response to these prototypical rhythmic episodes may be stronger as compared to similar maternal-infant cues that do not contain a social component. Although such comparison has not yet been tested, it is reasonable to assume that since maternal care is a time consuming, metabolically costly endeavor, bearing a critical impact on species continuity, the mother’s brain would not activate to its full capacity when resources are needed for other tasks but would cohere into its full expression to sustain these brief social moments.

With regards to *brain-behavior coupling*, studies have shown greater activations in areas of the maternal brain to naturalistic interactions that contain more social synchrony, both own infant (Atzil et al., 2011) and age-matched infant (Atzil et al., 2014). Specifically, levels of activations in subcortical dopamine-rich structures (VTA, nucleus accumbens) and areas of the mesolimbic dopaminergic pathways, particularly the ACC, correlated with the degree of observed synchrony. These findings suggest that links between maternal behavior and activations of the human maternal brain, particularly in dopamine-rich areas, may be specific to moments of synchrony.

As to *oxytocin sensitivity*, we hypothesize that mother’s neural response to social cues would be more impacted by OT administration compared to non-social cues. OT is an important modulator of the social brain (Zink and Meyer-Lindenberg, 2012) and underpins the reorganization of the maternal brain upon birth (Insel and Young, 2001). OT plays a critical role in neural plasticity at the molecular and network assembly level and such plasticity is required to augment the salience and reward value of the infant to its mother (Marlin et al., 2015; Oettl et al., 2016; Valtcheva and Froemke, 2019) and both experimental and knockout studies demonstrate the causal role of OT in the initiation of maternal behavior (Higashida et al., 2010; Lopatina et al., 2012). Human studies have shown that peripheral levels of OT are associated with sensitive and synchronous parenting (Feldman et al., 2011a), and activation of the parent’s brain is linked with parental OT levels (Abraham et al., 2014; Rilling and Mascaro, 2017). Intranasal oxytocin administration targets social moments by increasing the prototypical social behaviors in parent and child and elevating their coordination (Naber et al., 2010; Weisman et al., 2012a), and it is therefore hypothesized that OT effects would be specific to the social context. However, the literature is mixed on whether OT administration may increase or attenuate these socially-driven neural activations (Chen et al., 2017; Grace et al., 2018; Wang et al., 2017; Wigton et al., 2015). Indeed, while some studies show increased activation of limbic, paralimbic, and temporal structures to oxytocin administration, consistent with the “social salience” hypothesis of oxytocin (Shamay-Tsoory and Abu-Akel, 2016), others demonstrated attenuation of these same regions to OT administration, consistent with the anxyolitic model on OT (Neumann and Slattery, 2016). As the current consensus is that the effects of OT are time-, person- and context-sensitive (Bartz et al., 2011), the direction of its effect on the maternal neural response to social stimuli remains an open question.

Finally, we explored the existence of “*temporal engrams*” in the maternal brain in response to the prototypical repetitive-rhythmic social moments. Engrams describe the distinct neural processes by which the attachment target is “imprinted” in the brain (Walum and Young, 2018). In humans, the experience of social synchrony and its specific rhythms may function to build and amplify temporal patterns in the brain. The brain’s sensitivity to temporal patterns, the “sociotemporal brain” (Schirmer et al., 2016), is thought to rely particularly on insular and temporal regions that gauge the durations and patterns of social stimuli and ground them in the here-and-now (Craig, 2009; Seth and Friston, 2016). Temporal patterns in the brain are suggested to be impacted by interoceptive processes, which provide the foundation for the sense of self based on representations of bodily signals that form predictions (“priors”) based on repeated experiences to enable the perception and attribution of meaning to incoming information (Allen and Tsakiris, 2018; Park et al., 2016; Salomon et al., 2016; Seth, 2013). Insular and temporal cortices are central to the brain’s perception of temporal regularities (Salomon et al., 2016; Schirmer et al., 2016; Wittmann, 2013) and have been implicated in brain-to-brain synchrony during social interactions (Anders et al., 2011; Hasson et al., 2004; Kinreich et al., 2017; Nummenmaa et al., 2012). As an open-ended research question, we thus explored whether, when mother’s brain is repeatedly imaged, insular and temporal cortices would show a consistent pattern of activation, a *temporal engram* to attachment stimuli. However, we expected that only the special synchronous social moments would elicit this unique temporal pattern, which would be found in the AI and structures of the temporal cortex (e.g., STS, TP).

As such, the goal of the current study was to describe the neural signature of social cues in the human mother’s brain. To test this, we expanded on a well-researched paradigm into the parental brain, which involves presentation of individually-tailored stimuli collected in the home ecology (Atzil et al., 2011; Elmadih et al., 2016; Noriuchi et al., 2008) and included three separate conditions that depict typical mother-child social and non-social contexts in their home environment. Across these conditions, mothers were filmed sitting next to their child in the same level and distance, to control for differences in physical proximity and posture. In the first condition mothers sat next to their infant while being otherwise engaged (Condition I, *Unavailable*); in the second, mothers sat facing the infant but did not engage in social interactions (Condition II, *Unresponsive*); in the third, mothers engaged in a prototypical, rhythmic social play of pick-a-boo (Condition III: *Social*). Mothers were imaged twice in a double-blind within-subject placebo-control design and observed the same three conditions viewing themselves (Self) and an unfamiliar mother-infant dyad (Other) infant once following the administration of oxytocin (OT) and once after placebo (PBO).

We expected to find the four distinctly-social processes in response to the social condition, but not to the other conditions. First, the social condition would elicit the greatest response across the maternal caregiving network compared to the other conditions. Next, only activations to the social condition would show brain-behavior coupling with mother-infant synchrony assessed in the home environment, but no such brain-behavior links would emerge with the non-social conditions. Third, we expected that OT would impact specifically the mother’s brain response to the social condition, and tested whether these OT-mediated neural responses would follow the “social salience” hypothesis (i.e., increased brain activations to social condition under OT) or the anxiolytic model of OT (decreased brain activation under OT). Fourth, we explored whether the temporal pattern of insula and structures of the temporal cortex (STS, TP) activation to the social condition, but not to the non-social condition, would show higher consistency between the two imaging sessions (“temporal engrams”). Finally, mother’s brain activations would show a differential response to her own, as compared to an unfamiliar infant and that these would increase in the social condition.

## Results

### Salivary oxytocin following administration and micro-level synchrony across conditions

To validate our procedure, we first examined whether micro-level synchrony was indeed higher in the social compared to the unavailable and unresponsive conditions, to ascertain that this condition exposed mothers to high levels of synchrony. As expected, a repeated measures ANOVA [F(1.16,25.56)= 49.16, p< .001] revealed that stimuli in the *Social* condition comprised significantly more synchrony (Mean=5.96, SD=3.87) compared to the *Unavailable* (Mean=0.304, SD=1.26) and *Unresponsive* (Mean=0.00, SD=0.00) conditions (Figure 1), validating our paradigm.

**Figure. 1.**
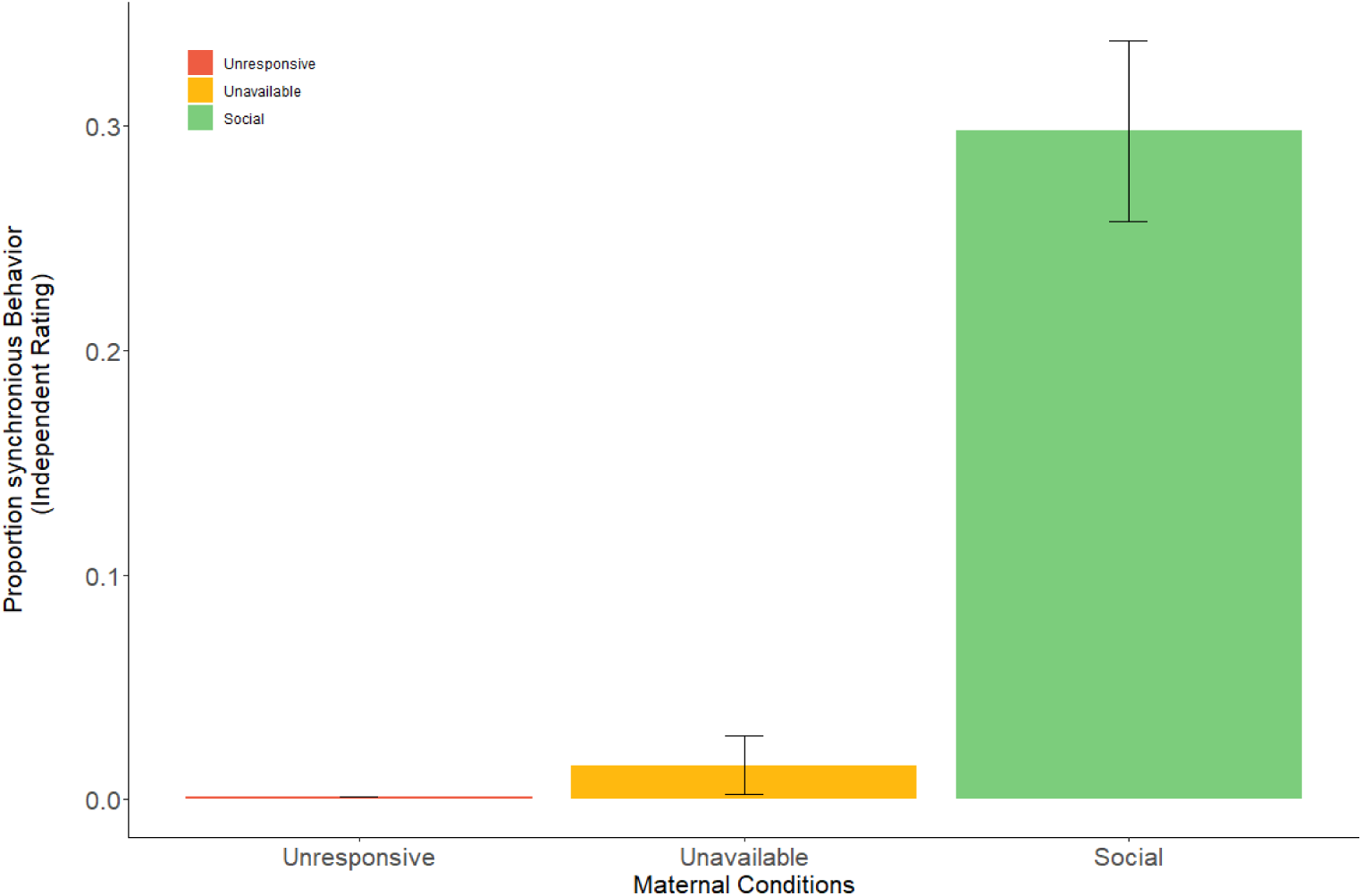
Proportion of mother-infant synchronous behavior in the three maternal conditions. Mother-infant synchrony occurred more during face-to-face interaction in the social maternal condition compared to the unavailable and unresponsive maternal conditions. All effects were Greenhouse-Geisser corrected. Error bars represent standard error of the mean.

Next, to validate the OT manipulation, we tested whether peripherally-measured OT levels were indeed higher after OT administration. A 2×3 (*PBO-OT* × *Time*) repeated measures ANOVA on mother salivary oxytocin levels (pg/mL) showed a significant *PBO-OT* × *Time* interaction effect [F(1.1,24.09)= 10.01, p < .01], *PBO-OT* main effect [F(1,22)= 10.46, p < .01] and *Time* main effect [F(1.09,23.86)=11.03, p<.01]. As expected, following OT administration mothers showed a marked increase in OT levels (Mean=826.75, SD=1135.25) compared to the baseline (Mean=21.49, SD=13.61) and to the recovery samples (Mean=193.86, SD=303.56). In contrast, no significant increase in peripheral OT was observed following PBO administration (Figure supplement 1).

### fMRI Whole Brain analysis

To examine brain regions associated with our conditions, a whole-brain 3 factorial ANOVA (*Maternal Condition* × *Self-Other*× *PBO-OT*) was calculated within BrainVoyager software. The analysis revealed a significant, FDR corrected, *Maternal Condition* main effect. A 200 voxels cluster size was used to extract volumes of interest (VOIs) from all regions that demonstrated significantly differential activity. The ANOVA revealed a widespread network of activations across the insula, superior-frontal and temporal areas in the cortex. Regions showing differential activations across the three maternal conditions included the cingulate gyrus, left insula, bilateral frontal lobe areas, bilateral STG to TP, bilateral parahippocampal gyrus, bilateral anterior cerebellum, bilateral globus pallidus-Putamen, right MOG, and right Cuneus (See Table 1, Figure 2).

**Table 1.**
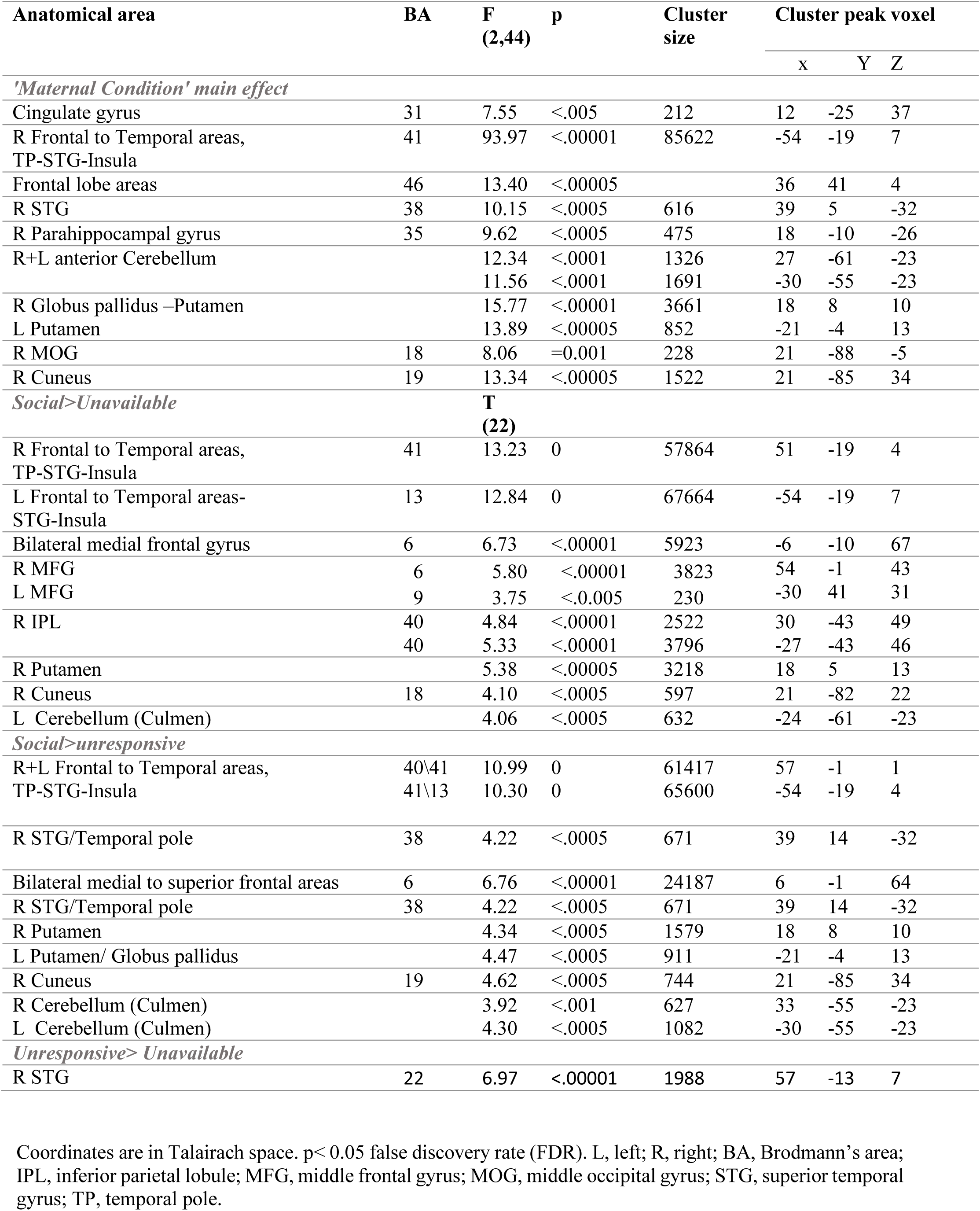
Coordinates of activation peaks (whole brain ANOVA results)

| Anatomical area | BA | F<br>(2,44) | p | Cluster<br>size | Cluster peak voxel |  |  |
| --- | --- | --- | --- | --- | --- | --- | --- |
|  |  |  |  |  | x | Y | Z |
| 'Maternal Condition' main effect |  |  |  |  |  |  |  |
| Cingulate gyrus | 31 | 7.55 | <.005 | 212 | 12 | -25 | 37 |
| R Frontal to Temporal areas,<br>TP-STG-Insula | 41 | 93.97 | <.00001 | 85622 | -54 | -19 | 7 |
| Frontal lobe areas | 46 | 13.40 | <.00005 |  | 36 | 41 | 4 |
| R STG | 38 | 10.15 | <.0005 | 616 | 39 | 5 | -32 |
| R Parahippocampal gyrus | 35 | 9.62 | <.0005 | 475 | 18 | -10 | -26 |
| R+L anterior Cerebellum |  | 12.34 | <.0001 | 1326 | 27 | -61 | -23 |
|  |  | 11.56 | <.0001 | 1691 | -30 | -55 | -23 |
| R Globus pallidus –Putamen |  | 15.77 | <.00001 | 3661 | 18 | 8 | 10 |
| L Putamen |  | 13.89 | <.00005 | 852 | -21 | -4 | 13 |
| R MOG | 18 | 8.06 | =0.001 | 228 | 21 | -88 | -5 |
| R Cuneus | 19 | 13.34 | <.00005 | 1522 | 21 | -85 | 34 |
| Social>Unavailable |  | T<br>(22) |  |  |  |  |  |
| R Frontal to Temporal areas,<br>TP-STG-Insula | 41 | 13.23 | 0 | 57864 | 51 | -19 | 4 |
| L Frontal to Temporal areas-STG-Insula | 13 | 12.84 | 0 | 67664 | -54 | -19 | 7 |
| Bilateral medial frontal gyrus | 6 | 6.73 | <.00001 | 5923 | -6 | -10 | 67 |
| R MFG | 6 | 5.80 | <.00001 | 3823 | 54 | -1 | 43 |
| L MFG | 9 | 3.75 | <.0.005 | 230 | -30 | 41 | 31 |
| R IPL | 40 | 4.84 | <.00001 | 2522 | 30 | -43 | 49 |
|  | 40 | 5.33 | <.00001 | 3796 | -27 | -43 | 46 |
| R Putamen |  | 5.38 | <.00005 | 3218 | 18 | 5 | 13 |
| R Cuneus | 18 | 4.10 | <.0005 | 597 | 21 | -82 | 22 |
| L Cerebellum (Culmen) |  | 4.06 | <.0005 | 632 | -24 | -61 | -23 |
| Social>unresponsive |  |  |  |  |  |  |  |
| R+L Frontal to Temporal areas,<br>TP-STG-Insula | 40\41 | 10.99 | 0 | 61417 | 57 | -1 | 1 |
|  | 41\13 | 10.30 | 0 | 65600 | -54 | -19 | 4 |
| R STG/Temporal pole | 38 | 4.22 | <.0005 | 671 | 39 | 14 | -32 |
| Bilateral medial to superior frontal areas | 6 | 6.76 | <.00001 | 24187 | 6 | -1 | 64 |
| R STG/Temporal pole | 38 | 4.22 | <.0005 | 671 | 39 | 14 | -32 |
| R Putamen |  | 4.34 | <.0005 | 1579 | 18 | 8 | 10 |
| L Putamen/ Globus pallidus |  | 4.47 | <.0005 | 911 | -21 | -4 | 13 |
| R Cuneus | 19 | 4.62 | <.0005 | 744 | 21 | -85 | 34 |
| R Cerebellum (Culmen) |  | 3.92 | <.001 | 627 | 33 | -55 | -23 |
| L Cerebellum (Culmen) |  | 4.30 | <.0005 | 1082 | -30 | -55 | -23 |
| Unresponsive> Unavailable |  |  |  |  |  |  |  |
| R STG | 22 | 6.97 | <.00001 | 1988 | 57 | -13 | 7 |

Coordinates are in Talairach space. $p < 0.05$ false discovery rate (FDR). L, left; R, right; BA, Brodmann's area; IPL, inferior parietal lobule; MFG, middle frontal gyrus; MOG, middle occipital gyrus; STG, superior temporal gyrus; TP, temporal pole.

**Figure 2.**
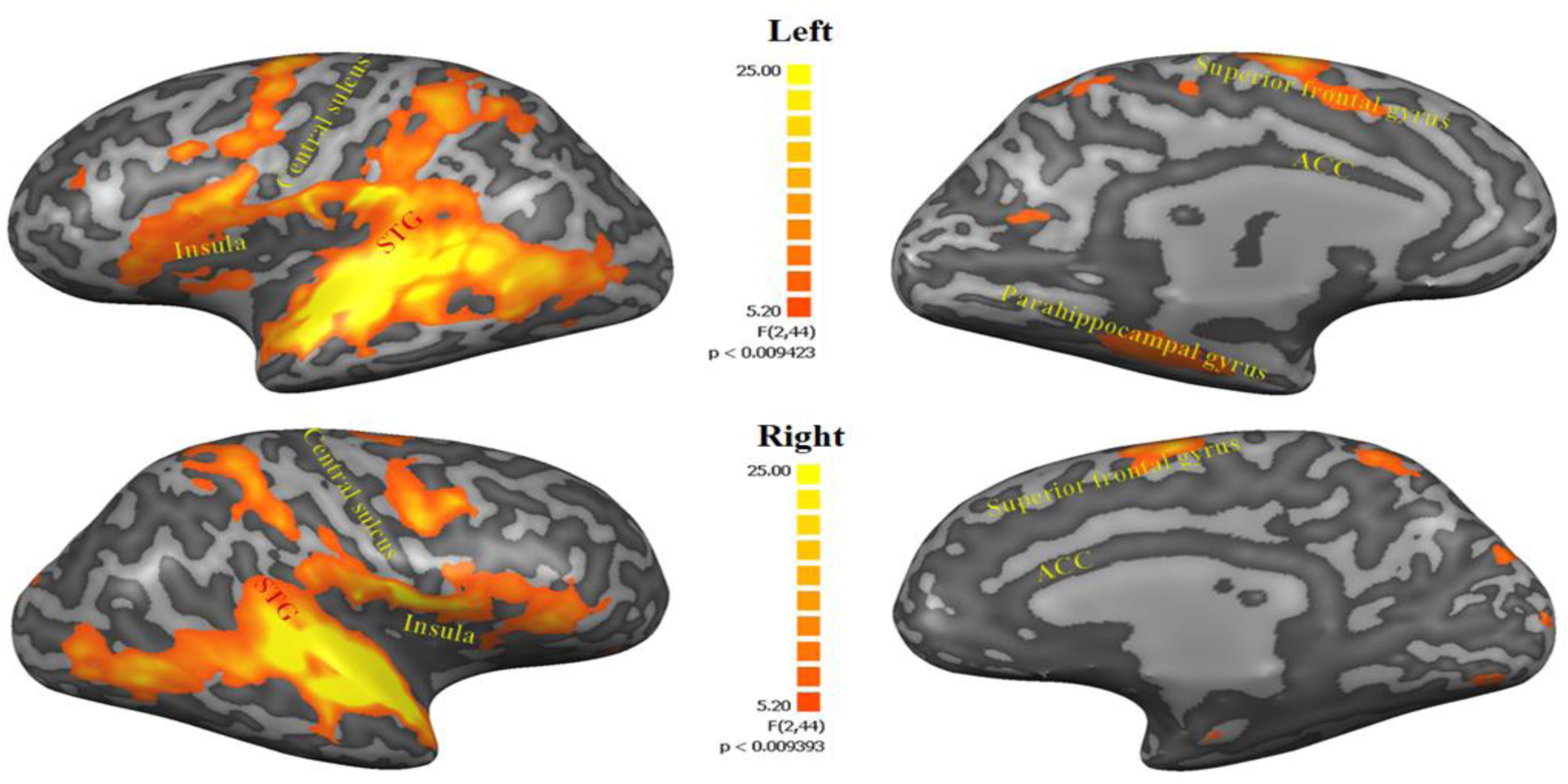
Maternal Condition main effect. Figures representing regions within 3 factorial ANOVA ‘maternal condition’ main effect (FDR corrected, Cluster threshold of 200) including the cingulate gyrus, left insula, bilateral frontal lobe areas, bilateral STG toTP, bilateral parahippocampal gyrus, bilateral anterior cerebellum, bilateral globus pallidus-Putamen, right MOG, right cuneus. STG, superior temporal gyrus; TP, temporal pole; MOG, middle occipital gyrus.

To examine the origin of the maternal condition main effect, three planned contrasts were used: *Social* (Self+Other+OT+PBO) > *Unavailable* (Self+Other+ OT+PBO) (Figure 3.A) and *Social* (Self+Other+OT+PBO)> *Unresponsive* (Self+Other+OT+PBO) (Figure 3.B). As seen in Figure 3, both contrasts elicited activations in the temporal and frontal cortices including the STG to TP, the insula, and the medial frontal gyrus, in addition to activations in subcortical structures in the basal ganglia (the putamen and the globus pallidus) and in the cerebellum, which were significantly higher in the *Social* compared to the other conditions, supporting our first hypothesis. The contrast of *Unresponsive* (Self+Other+OT+PBO) > *Unavailable* (Self+Other+ OT+PBO) was examined as well. No significant, FDR corrected, results were found for *Self-Other* or *PBO-OT* main effects. All activations for all contrasts can be seen in Table 1.

**Figure 3.**
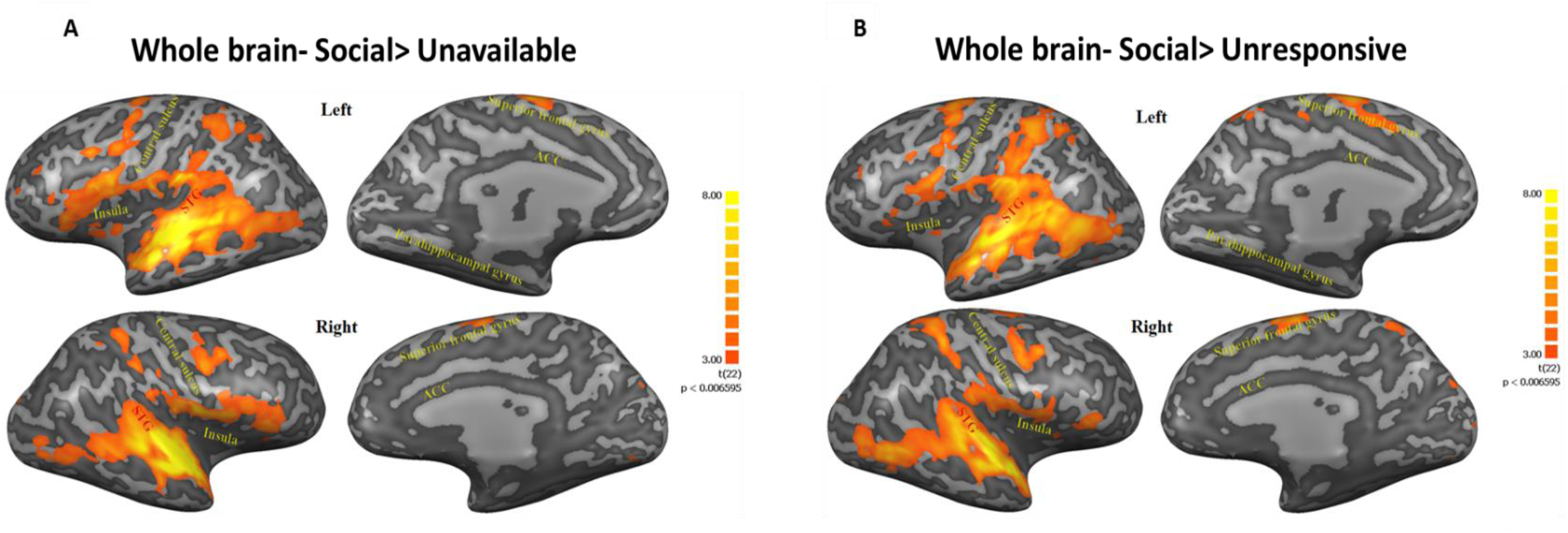
**A, B.** Figures represent regions within post hoc contrasts (Social>Unavailable; Social>Unresponsive respectively) conducted to further examine the significant maternal condition main effect found in the whole brain 3 factorial ANOVA. Note that similar areas were elicited in both contrasts. This highlights the extensive activity along regions ranging from the insula, STG to TP and areas in the frontal cortex under the social condition. Subcortical structures in the basal ganglia and the cerebellum were also activated in both contrasts. Results are FDR corrected with Cluster threshold of 200. Brain regions are defined in table 1. STG, superior temporal gyrus; TP, temporal pole.

### Brain-Behavior coupling

To test out second hypothesis, which maintains that brain–behavior coupling is specific to the Social condition, we computed Pearson’s correlations between ROIs activation to the *Self-Social* condition under placebo (representing the mother’s brain response under natural circumstances), and mother-infant synchrony measured during free-play interaction in the home ecology. Mother-Infant synchrony correlated with activity in the bilateral insula, ACC, and VTA, indicating that more synchronous mothers exhibited greater activation in these areas to videos depicting themselves interacting with their infants (Figure 4). Mother infant synchrony was also positively correlated with activity in the VTA (r_p_=0.461, p=0.03). No significant correlations emerged between mother-infant synchrony and activation of any ROI in the *Unavailable* or *Unresponsive* conditions. Under OT, correlations were non-significant.

**Figure 4.**
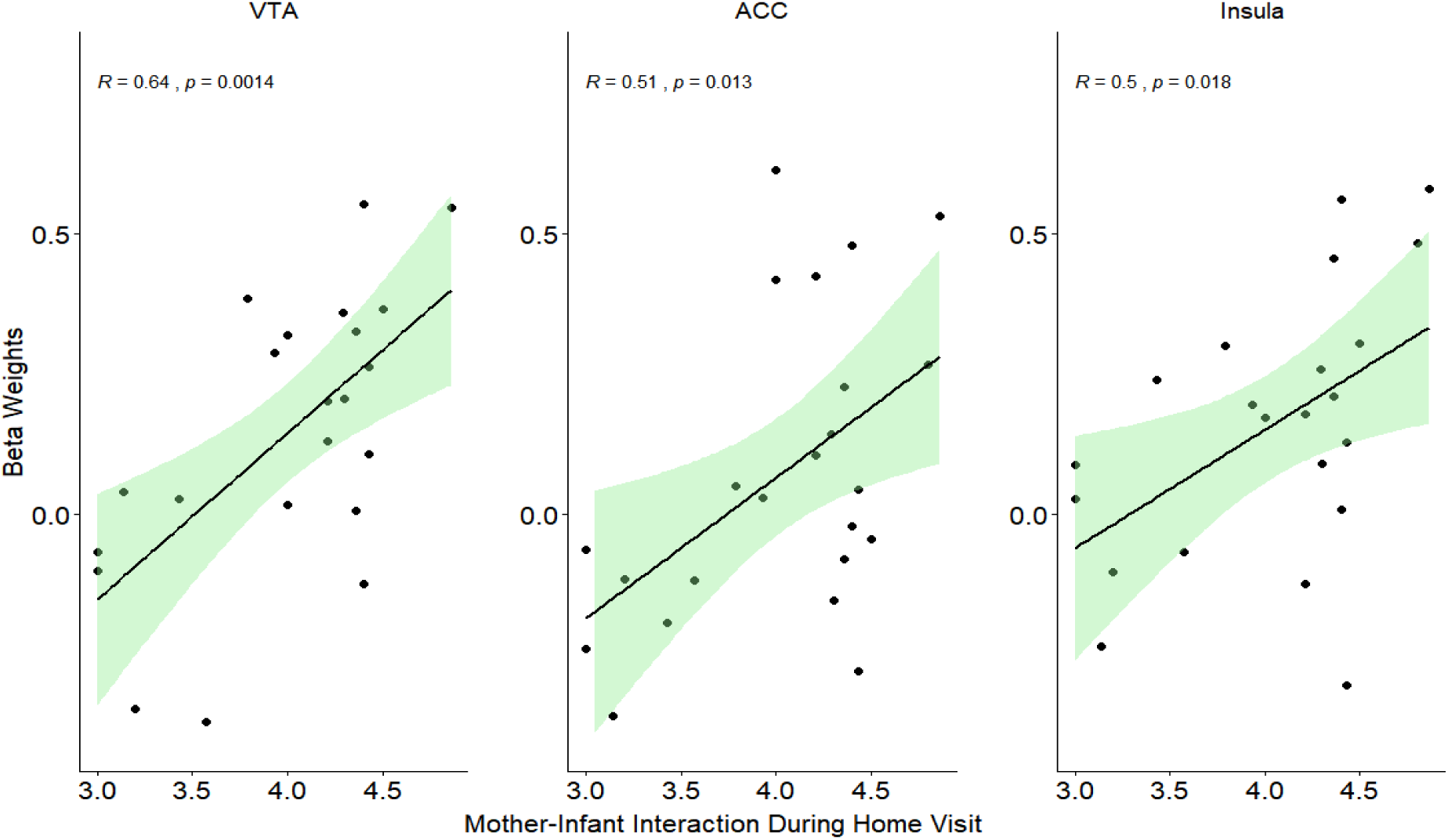
Regression lines of significant correlations between mother-infant synchrony during a free-play interaction at home visit and activation in the VTA, insula and ACC under ‘Social’ maternal condition and placebo. VTA, ventral tegmental area; ACC, anterior cingulate.

### Oxytocin effects on ROIs activation

In order to examine the eight preregistered regions of interest, a 3 factorial repeated measures ANOVA (*Maternal Condition*× *Self-Other*× *PBO-OT*) was performed on the Beta values extracted from each of the ROIs. No three-way interaction was found in any ROI (see Table 2).

**Table 2.**
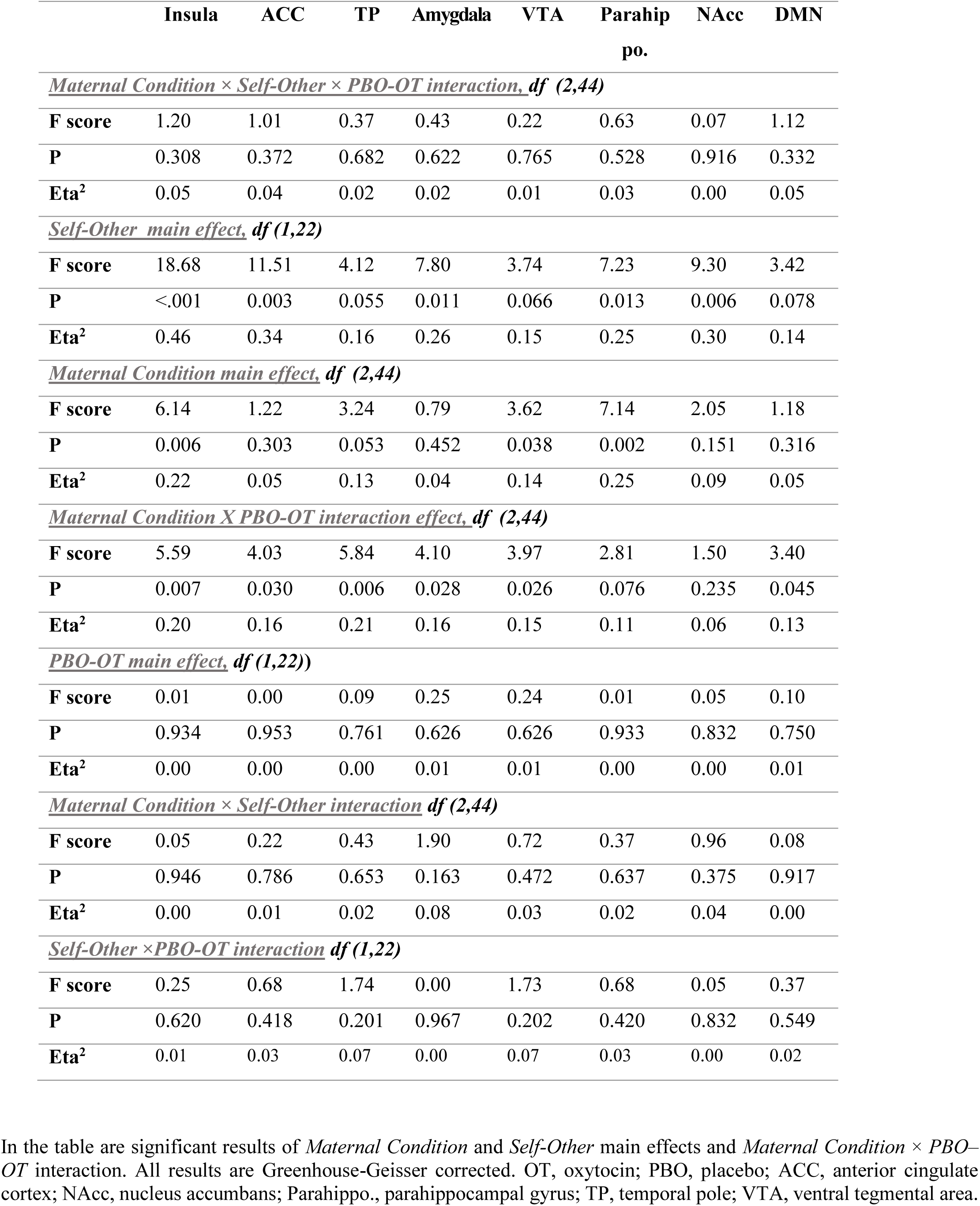
Results of repeated measures ANOVA (*Maternal Condition* × *Self-Other* × *PBO- OT*) for preregistered ROIs.

Significant *Self-Other* main effect was found in five ROIs (Figure supplement 2). Greater activation in response to the Self compared to Other-stimuli was found in the insula (Mean_self_=0.04, SD_self_=0.13; Mean_other_=-0.08, SD_other_=0.17) and the ACC (Mean_self_=-0.08, SD_self_=0.18; Mean_other_=-0.21, SD_other_=0.21). In contrast the amygdala (Mean_self_=0.10, SD_self_=0.17; Mean_other_=0.23, SD_other_=0.18), parahippocampal gyrus (Mean_self_=-0.03, SD_self_=0.17; Mean_other_=0.05, SD_other_=0.13) and the NAcc (Mean_self_=-0.17, SD_self_=0.27; Mean_other_=0.00, SD_other_=0.21) were more activated during the Unfamiliar-Other compared to the Self-stimuli (Table 2).

Main effect for maternal condition was found in several of the preregistered ROIs. In the insula it was driven by high responses to social (Mean=0.06, SD=0.18) compare to unavailable (Mean=-0.04, SD=0.16) and to unresponsive (Mean=-0.07, SD=0.16). In the VTA it was driven by high responses to unavailable conditions (Mean=0.03, SD=0.12) compared to unresponsive (Mean=-0.043, SD=0.159). In the parahippocampal gyrus it was driven by a high response to unavailable (Mean=0.09, SD=0.13) compare to social (Mean=-0.05, SD=0.20). No significant main effect for PBO-OT was found in any of our ROIs.

Critically, our main finding was defined by an interaction effect of *Condition* × *PBO-OT*, observed in 6 out of 8 preregistered ROIs (all results are shown in Figure.5, Table 2). In five ROIs, including the insula, amygdala, ACC, STG, and VTA, one way ANOVA conducted to PBO and OT separately, revealed that the interaction was driven by significant differences between the three conditions under placebo, while under oxytocin no such differences were found. In the DMN, significant differences between conditions emerged under OT and not under placebo. For other regions that did not show significant effects see table 2.

**Figure 5.**
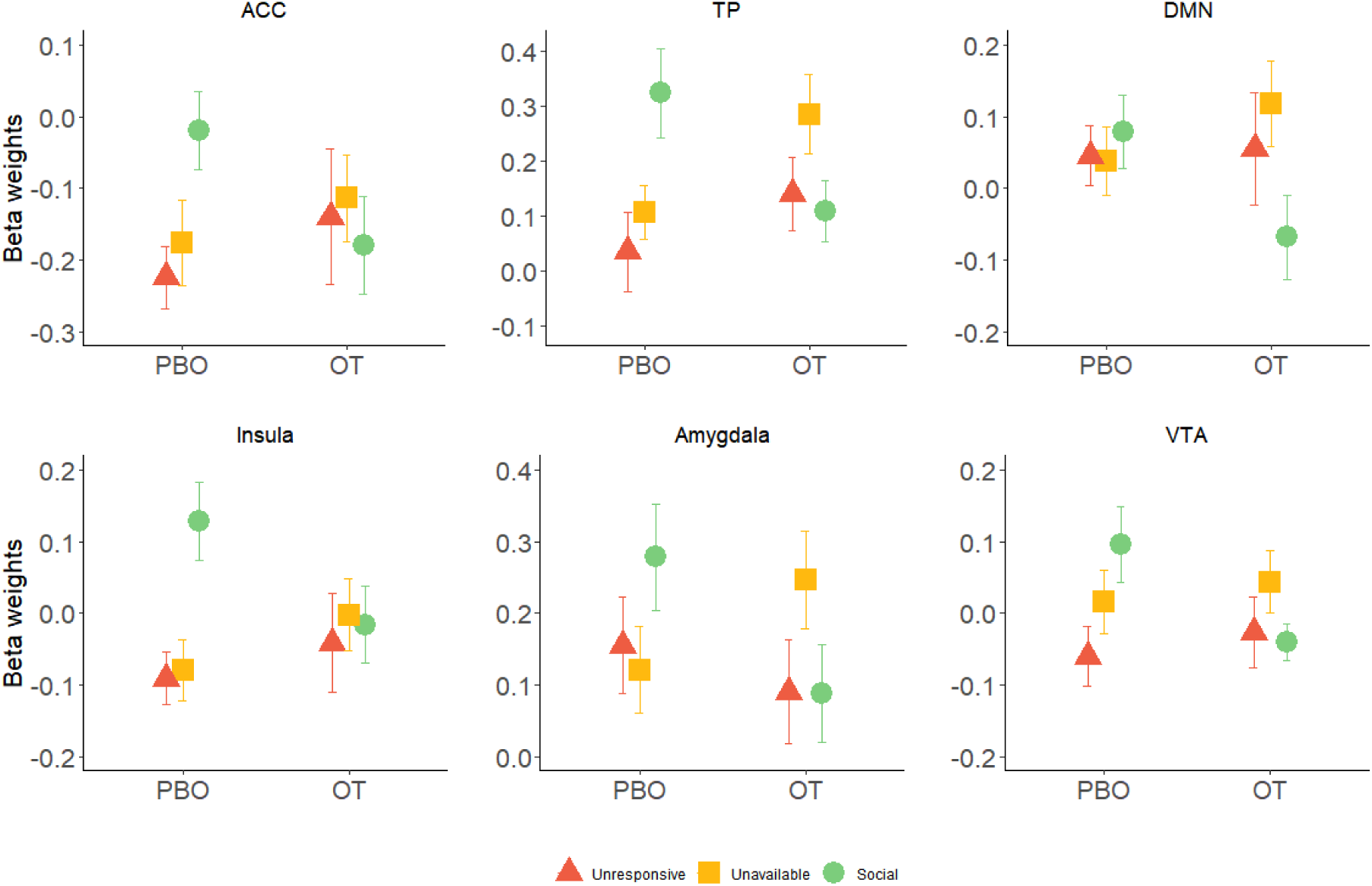
Significant interaction effects of *Maternal Condition × PBO-OT in 6 preregistered ROIs*. In the ACC, insula, TP, amygdala and VTA differences between brain response to the maternal conditions were observed under placebo (PBO), with highest activation in response to the social condition (in green). After oxytocin (OT) administration these differences disappeared. In the DMN the opposite pattern was observed: under oxytocin there were differences and under placebo not. Bars depict Standard error of the mean. PBO, placebo; OT, Oxytocin; ACC, Anterior cingulate; DMN, Default mode network; TP, Temporal pole; VTA, ventral tegmental area.

### Within subject correlation (WSC)

Finally, we wished to explore the temporal consistency of activation patterns in our ROIs across the two scans for the three maternal conditions. For this end, we used Within Subject Correlations (WSC) for each participant between the oxytocin and placebo scans and for each of the three maternal conditions. A two factorial repeated measures ANOVA (ROI × *Maternal Condition*) revealed a significant main effect of *Condition* [F (1.97,43.29)=5.08, p=0.01]. Post hoc comparisons revealed significantly stronger WSC under the *Social* condition (Mean=0.14, SD=0.05) compared to the *Unresponsive* condition (Mean=0.03, SD=0.02). This effect was driven by the insula [t(22)=2.60, p<0.05] (Mean_social_=0.20, SD_social_=0.23; Mean_unresponsive_=0.05, SD_unresponsive_=0.17); TP [t(22)=2.58, p<0.05] (Mean_social_=0.23, SD_social_=0.25; Mean_unresponsive_=0.04, SD_unresponsive_=0.21), and amygdala [t(22)=1.83, one tailed p<0.05] (Mean_social_=0.19, SD_social_=0.17; Mean_unresponsive_=0.07, SD_unresponsive_=0.25), which showed significant differences between WSC for condition (Figures 6.A-E).

**Figure 6.**
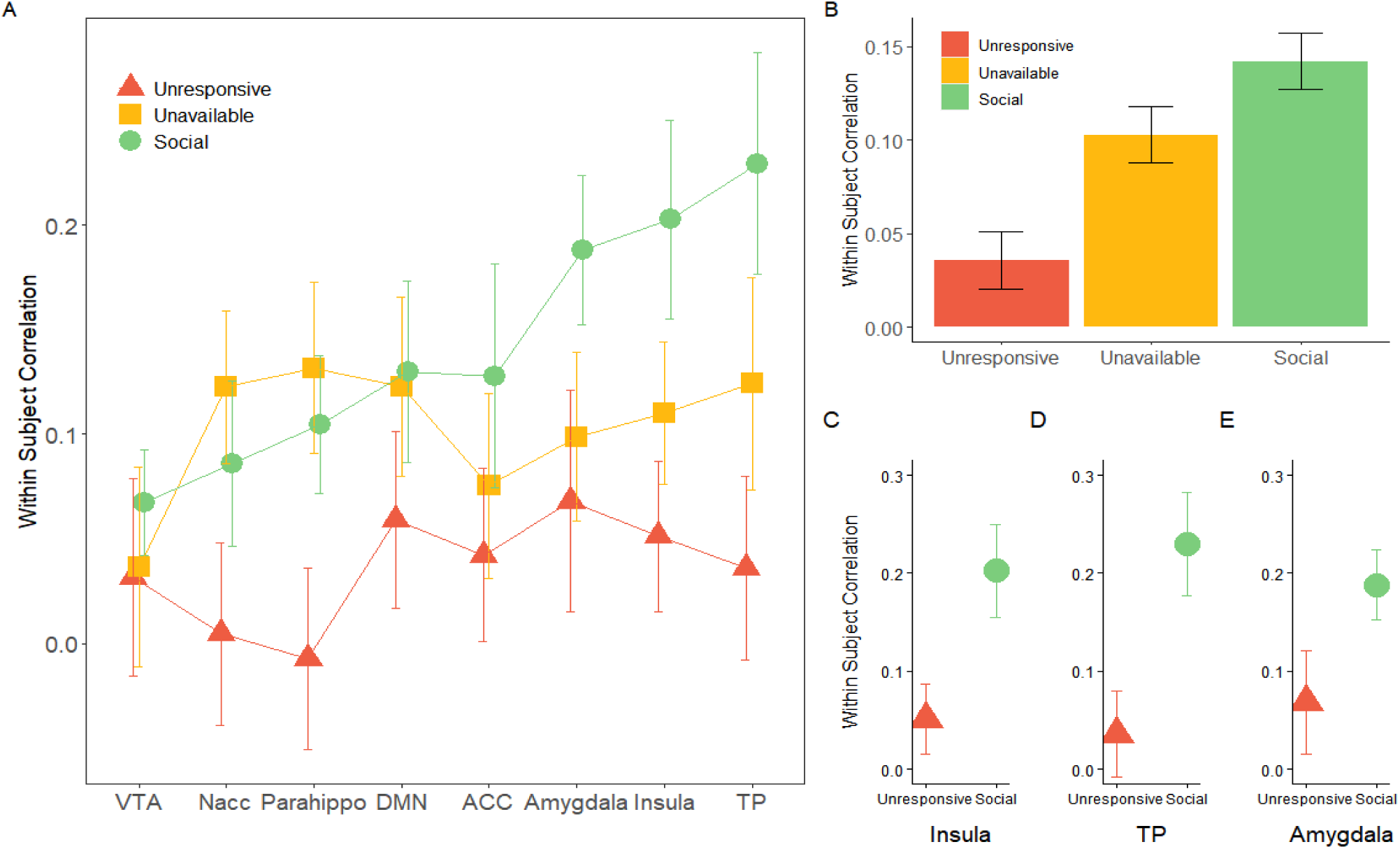
Within subject correlation (WSC) across ROIs: WSC represent the temporal activation pattern consistency of the ROIs under the three maternal conditions. It was calculated for each subject by Pearson’s correlation for the BOLD time course of each condition between the OT and PBO runs. **A.** 8×3 repeated measures ANOVA (*ROI* × *Maternal Condition*) revealed a significant main effect of *Maternal Condition* **B.** Under the *social* maternal condition, the WSC was higher compared to the *unresponsive* maternal condition. **C, D, E.** This effect was pronounced and significant in the insula, TP and amygdala. Bars depict Standard error of the mean. OT, oxytocin; PBO, placebo; TP, temporal pole.

## Discussion

For social mammals born with an immature brain who require the external regulation of the mother’s mature brain for development, moments of maternal-infant social contact hold a special significance. These brief social moments integrate multiple well-orchestrated bio-behavioral processes that organize the maternal brain, trigger the species-typical caregiving behavior, and carry imprinting-like effect on the infant’s brain. Across mammalian species, the cross-generation transmission of sociality originates in the consolidation of the “maternal brain” from which, through caregiving behavior, a similar network is sculpted in the infant (Numan, 2020). Our study is the first to assess the maternal neural response to these social moments, versus other moments of mother-infant presence, in order to shed further light on the cross-generation transmission of human sociality.

To tap the mother’s neural response to social interactions, we employed a double-blind OT/placebo crossover design; hence, our study is the first to image the postpartum brain twice within a brief time-span in response to multiple ecological contexts. Results describe four processes by which social moments make their mark on the maternal brain. First, we found that under typical conditions (i.e. PBO), areas of the “maternal caregiving network”, including both the mammalian-general and human-specific structures, showed elevated activity in response to the social, compared to the other conditions. Second, brain-behavior coupling was found only to the maternal socially-driven neural response, underscoring the sensitivity of these activations to actual social patterns in the natural habitat. Third, we show that the effects of OT on the maternal brain were more notable during the social condition and OT decreased neural activation to social stimuli, indicating that neural response to social moments is more sensitive to OT. Finally, only during the social condition we found consistency in patterns of activation across repeated imaging in insula and TP, areas of the “sociotemporal brain” (Schirmer et al., 2016). Overall, our findings highlight the importance of these social moments for the maternal brain. However, in contrast to our expectation, we found no interaction between condition and own-unfamiliar infant, indicating that the neural response to social moments did not differentiate own from other infant. This may suggest the salience of these highly-arousing, species-typical universal social exchanges is especially high and may trump differentiation of self and other, but further research is needed to fully understand these findings.

Brain-behavior coupling was found between the degree of mother-infant synchrony observed in the home ecology and mother’s brain activation in the VTA, ACC, and insula and activations in the VTA responded to the degree of synchrony to both own and unfamiliar infant. The VTA and ACC are part of the mesolimbic dopamine pathway and we have previously shown that when naturalistic interactions contain more social synchrony, both the mother’s own and unfamiliar infant, they correlate with greater activations of the dopaminergic pathway (Atzil et al., 2011). The VTA plays a key role in mammalian maternal care and studies in animal models show that the formation of new projections from the hypothalamic MPOA to the VTA, primed by the OT increase during pregnancy and childbirth, renders the infant the most rewarding stimulus to its mother and this stimulates the mother’s approach orientation and increases offspring survival (Numan, 2020). The insula plays a key role in the maternal brain, enabling the mother to resonate with the infant’s state online and to provide an overall allostatic function for the infant’s emerging bodily representations.

Our third process proposed special sensitivity of socially-driven neural activations to OT administration. Overall, we found a similar pattern of results across five areas, including the amygdala, VTA, insula, ACC, and TP. While these regions showed increased activity to the social context under natural conditions (i.e. PBO), OT suppressed these social-specific activations and leveled out the mother’s neural response so that under OT, no differences were found between the social and non-social conditions. A recent study assessing the effects of OT administration via both intravenous and intranasal pathways on neural response (resting state) showed that OT via both pathways decreased activity in the amygdala, insula, parahippocampal gyrus, and TP, and the decrease was mediated by elevations in peripheral OT levels (Martins et al., 2020). Our findings similarly show a combination of increase in peripheral OT levels (Figure supplement 2) and the current findings showing muting of the neural response to social interactions in amygdala, insula, and TP extend previous findings in resting state to the neural processing of social stimuli. In combination, these findings may suggest that one pathway s for the anxiolytic effects of OT may be via the attenuation of activity in a limbic network that monitors salience cues, gauges danger signals, and integrates exeroceptive and interoceptive inputs to give immediacy and focus to ongoing events.

OT plays an important role in tuning the maternal brain and in triggering the cross-generational transfer of sociality. Humans are wired for social behavior via activity of the mammalian caregiving network, which contains abundant OT receptors (Sokolowski and Corbin, 2012). Initiation of maternal care in rodents involves a two-step processed; first, OT leads to long-term depression in amygdala to suppress social avoidance to infant stimuli (Gur et al., 2014), and next, OT connects with dopamine through striatal neurons that encode for both OT and dopamine D1 receptors (Olazábal and Young, 2006). This enables dopamine neurons that are particularly sensitive to identify sensory-motor general reward patterns to also encode the temporal patterns of *social* reward (Báez-Mendoza and Schultz, 2013; Ross and Young, 2009). Such process underpins the formation of maternal-infant bonding and allows the brain to internalize the social partner and its preferences, encode relationship-specific socio-temporal patterns, and draw reward from the matching of social actions of self and partner, that is, interactive synchrony (Báez-Mendoza and Schultz, 2013; Schultz, 2016). The tight cross-talk of oxytocin and dopamine triggered by maternal care enables the mother’s brain to form a neural integration of sensory experiences from the infant’s smell, touch, babbling, and cute face, and integrate them into an overall representation that contains the dyad-specific temporal patterns which underpins the attachment bond (Ross et al., 2009).

Yet, while OT constructs the temporal patterns of attachment in the maternal brain through connectivity with DA neurons that are time-sensitive and reward-focused and directs these patterns to the exclusivity of the mother-infant bond and its unique social rhythms, OT also has well-known anxiolytic properties (Neumann, 2008). The soothing function of OT is critical for survival. During labor, OT surges to levels substantially higher than daily levels and such high OT concentrations sooth the mother’s pain and stress through the regulatory effects of OT on the hypothalamic-pituitary-adrenal-axis and on sympathetic activity (Carter, 2014; Neumann and Slattery, 2016). OT enables tranquility during the birth process, attenuates amygdala response to external events to focus on the infant, and diminishes insular monitoring of unfamiliar interoceptive signals; lulling the mother’s brain from oscillating between extreme emotional states. Following OT administration, levels of OT rise well above their typical daily levels to levels unfamiliar to the body (Weisman et al., 2012b), and the anxiolytic properties of OT during the postpartum period, particularly in response to infant stimuli, may function in a similar way as during birth to sooth the mother’s brain and level-out the distinct response to different emotions.

In this context, it is interesting to note the response of the DMN to OT administration, which showed a somewhat different pattern of activations. We included the DMN in our preregistered ROIs in order to assess the “maternal caregiving network” in comparison with another well-characterized network known to be sensitive to self-related processing (Andrews-Hanna et al., 2014; Northoff et al., 2006; Peer et al., 2015) and to pinpoint the effects of social moments on the mother’s parenting-specific network. The DMN provides a useful comparison as it is thought to sustain the sense of self and is sensitive to self-other distinction across numerous types of stimuli (Davey et al., 2016; Salomon et al., 2014, 2009). Unlike the “maternal caregiving network” whose evolutionary role is to orient outward toward infant care and safety, the DMN is self-referential and inward-oriented. Here the DMN was selected as a unitary network (including the midline frontal and parietal, as well as lateral parietal nodes) to avoid multiple comparisons. The DMN showed no significant effect of self-versus other, likely due to the mother’s orientation towards external stimuli and possibly as a result of averaging the different regions of the DMN, which are known to have different functional selectivity to self-related stimuli (Araujo et al., 2015; Davey et al., 2016; Northoff et al., 2006; Salomon et al., 2014). However, while OT decreased activations in both the maternal caregiving network and the DMN to social stimuli, the overall pattern of response in the two networks was distinct. Under natural conditions (i.e. PBO), the DMN showed no differential response to the social, compared to non-social contexts; however, OT significantly decreased DMN activation to the social condition, resulting not in an attenuation of differences among conditions, but in significantly lower DMN response to the social conditions. Thus while the DMN is typically insensitive to differences captured by the distinct maternal conditions, activity related to the social condition was reduced under OT, possibly due to reductions in other “upstream” regions which show such selectivity.

The temporal engram, the consistent pattern of activation in the maternal brain, was found only in response to the social moments which were characterized by high behavioral synchrony. It appears that whereas the *magnitude* of activations to social interactions was attenuated by OT, the *temporal patterns* (engrams) remained constant in the insula and TP, areas central to the perception of temporal regularities and the duration and patterning of social stimuli (Schirmer et al., 2016). The insula plays a key role in interoception. Through insular activations mothers can represent not only their own bodily state, but also the infant’s bodily signals of hunger, fatigue, satiety, and pain in their own brain and form predictive regularities of the infant’s physical needs (Abraham et al., 2019). In parallel, the infant’s emerging awareness of own body and construction of primitive representations of self are initiated by maternal-infant interactions (Atzil et al., 2018; Fotopoulou and Tsakiris, 2017). Indeed, interoceptive activations in the parental brain shaped children’s later social competencies and insular activation in the child’s brain to attachment cues (Pratt et al., 2019, 2018).

The insular cortex also plays an important role in allostasis, allowing the mother have an allostatic role for the infant’s emerging neurobiological systems in order to predict and satisfy the infant’s physiological needs in a “just in time” fashion (Sterling, 2012). Allostasis describes the anticipatory regulatory process by which the brain, based on a hypothalamic clock that underpins rest-activity cycles in every tissue of the body, allocates resources to the various bodily functions before physiological systems become disregulated (Schulkin and Sterling, 2019). Within the mother-infant bond, allostasis allows the mother to satisfy the infant’s needs before they arise and this is among the main mechanisms by which the maternal mature brain externally regulates the infant’s immature brain across mammalian species (Feldman, 2020; Hofer, 1987). The insula serves as a center of allostasis; its connectivity to targets within the body increases attention to bodily cues and its connection to the hypothalamus and von Economo neurons, which allow for rapid communication, make the insula particularly sensitive to temporal regularities. Our findings are the first to show consistent temporal pattern in the maternal insula in response to social mother-infant stimuli and, combined with the findings that insular response is associated with patterns of behavioral synchrony, our findings highlight the insular cortex as a special interface for building the dyad-specific brain-behavior rhythms of the attachment relationship.

Characterization of the maternal brain, requires much further research in humans. Much further research is needed to understand disruptions in the maternal brain, particularly to its distinct response to social moments, under various high risk conditions. Studies show attenuation of neural activations in the maternal brain to moments of mother-infant interaction in conditions such as poverty (Kim et al., 2017) or depression (Kim et al., 2016) and demonstrate that these are then transferred to insensitivity of the child’s brain to attachment cues (Pratt et al., 2019). Thus, our findings on the processes that underpin the mother’s neural response to social moments should be studied across various high-risk conditions. Furthermore, our findings can provide the basis for specific interventions that target maternal neural response to social moments and interactive synchrony and intervention outcome can be tested post-treatment. The reorganization of the maternal brain in the postpartum plays a critical role in species continuity across mammals species and underpin the cross-generational transmission of human social heritage and much further research is needed to fully understand its mechanisms in health and pathology, compare these processes to those of the paternal brain, and understand how cultural practices and meaning systems function to shape the mother’s neural response to social cues.

## Materials and Methods

### Participants

The initial sample included thirty-five postpartum mothers who were recruited through advertisements in various parenting online forums. Following recruitment, mothers underwent a brief phone screening for MRI scanning and postpartum depression using the Edinburgh Postnatal Depression Scale (Cox et al., 1987). Cutoff for joining the study was EPDS score of 8 and below (score above 9 indicates minor depression). Next, mothers were invited to a psychiatric clinic to be tested by a psychiatrist prior to OT administration. During this visit, mothers were interviewed using the Structured Clinical Interview for the DSM-IV (SCID) to assess current and past psychiatric disorders. None of participants met criteria for a major or minor depressive episode during the perinatal period, 97% did not meet criteria for any diagnosable psychopathology, and 86% did not meet criteria for any diagnosable psychopathology disorder during their lifetime. All participants in the study were married, cohabitated with the infant’s father, were of middle-or upper-class socioeconomic status, and completed at least some college.

Of the 35 participants, 3 did not complete a single scan (one due to medical problems and two due to claustrophobia). After initial processing, data of additional 9 mothers were excluded due to excessive head movement artifacts (6 participants, movements≥3mm) or abnormal brain activity in response to visual stimuli (3 participants). The final sample included twenty-three mothers (mean age = 28.8 years, SD = 4.7; EPDS mean score=2.48, SD=2.66) of 4-8-month-old infants (mean age = 5.78 months, SD = 1.25) who underwent scanning twice (46 scans). The study was approved by the Bar-Ilan University’s IRB and by the Helsinki committee of the Sourasky medical center, Tel Aviv (Ethical approval no. 0161-14-TLV). All participants signed an informed consent. Subjects received a gift certificate of 700 NIS (∼200USD) for their participation in all four phases of the study (diagnosis, home visit, and two imaging sessions).

### Procedure

Following psychiatric diagnosis, the study included three sessions. In the first, families were visited at home, several episodes of mother–infant interactions were videotaped, and mothers completed self-report measures.

Several films were used as stimuli for the functional magnetic resonance imaging (fMRI) sessions. The videos depicted three typical situations distinguished by the amount of mother-infant social interaction and included: 1. *Unresponsive* condition (mother sitting next to the infant busy with her cellphone), 2. *Unavailable* condition (mother facing infant but not interacting socially), and 3. *Social* condition (mother engaging in a face-to-face pick-a-boo interaction). In addition, a five minutes free interaction between mother and infant was filmed for behavioral coding. Instructions were: play with your infant as you normally do. In all interactions mothers were instructed to sit next to their infants in the same distance and used standard toy and infant seat.

In the second and the third sessions, mothers participated in brain scanning at the Tel-Aviv Sourasky Medical Center. Mothers were instructed to avoid food intake and breastfeeding two hours before arrival. Before each scan mothers received 24 IU of oxytocin or placebo intranasally in a randomized, placebo-controlled, double-blind, two-period crossover design. During each session, salivary samples for oxytocin were collected at three time-points: immediately after consent and before OT or Placebo administration, following OT or Placebo administration and before participants were taken for the fMRI scan, and after the scan. While in the scanner, mothers were presented with vignettes of individually-tailored stimuli of own mother–infant interactions and with fixed control stimuli of unfamiliar mother and infant interactions. On average, 14 days elapsed between the two scans (SD=11.67, mode=7, median=7), that were both scheduled for the morning hours (07:30-12:00). Study procedure is presented in Figure 7.

**Figure 7.**
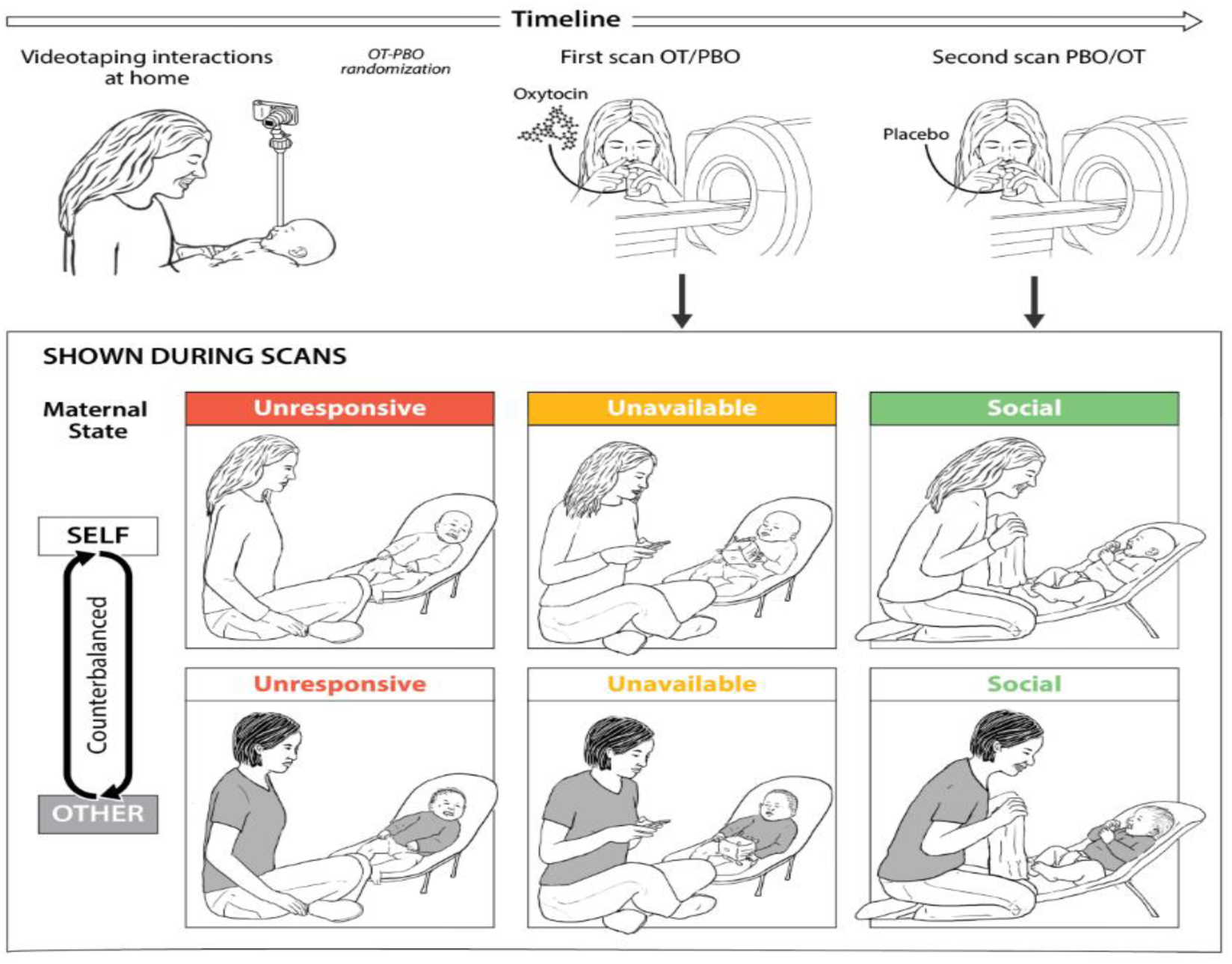
Research plan and fMRI paradigm. **A.** Experimental procedure. Mothers and infants were recruited 4-7 months post-partum and videotaped during a home-visit in the first session. Video vignettes of interactions were used as fMRI stimuli. During the second and third sessions mothers administered oxytocin or placebo before participating in brain scanning, in a randomized, placebo-controlled, double-blind, two-period crossover design. On average two weeks elapsed between scans. **B.** Experimental paradigm. Mothers were presented with six video vignettes of self and other (fixed control stimuli) mother-infant interactions depicting three maternal Conditions: Unresponsive-no interaction with the baby, mother does not respond to the baby (shown in red), Unavailable-minimal interaction, mother is busy, but respond to the baby when he/she signals (shown in yellow) and Social-mother and infant are engage in a pick-a-boo face-to-face social interaction (shown in green). Clips lasted 1 minute each and were previewed by rest with fixation period of 1 minute. A rest with fixation periods of alternately 15-18 seconds was presented between clips. Order of self-other and maternal conditions were counterbalanced between the two scans (PBO/OT).

### Oxytocin administration and salivary oxytocin collection and measurement

Mothers were asked to self-administer 24 IU of either oxytocin (Syntocinon Nasalspray, Novartis, Basel, Switzerland; three puffs per nostril, each containing 4 IU) or placebo. The placebo was custom-designed by a commercial compounding pharmacy to match drug solution without the active ingredient. The same type of standard pump-actuated nasal spray was used for both treatments. Three saliva samples were collected by passive drooling into a tube prior to inhaling oxytocin or placebo (baseline); 10-15 minutes after administration (post administration); and at the end of fMRI session (recovery). All samples were kept chilled and stored at −20°C. The concentration of OT was determined by Cayman-OT ELISA kit (Cayman Chemicals, Ann Arbor, Michigan, USA). Consistent with prior research we used ELISA (enzyme-linked immunosorbent assay), a method commonly used for analyzing hormones in saliva (Gordon et al., 2013; Rassovsky et al., 2019). In order to prepare the sample for measurement, samples underwent the following: 1. Freeze-thaw three cycles: freeze at −80°C and thaw at 4°C to precipitate the mucus; 2.

Centrifugations at 1500g (4000 rpm) for 30 minutes; and 3. The supernatant was transferred into clean tube, and stored at −20°C until assayed. Concentration of OT in these samples was determined according to the manufacturer’s kit instructions. The inter-assay coefficients of samples and controls were less than 18.7%, in the rage reported by the manufacture.

### MRI scans

#### Data acquisition

Magnetic Resonance Imaging (MRI) data was collected using a 3T scanner (SIEMENS MAGNETOM Prisma syngo MR D13D, Erlangen, Germany) located at the Tel Aviv Sourasky Medical Center. Scanning was conducted with a 20-channel head coil for parallel imaging. Head motion was minimized by padding the head with cushions, and participants were asked to lie still during the scan. High resolution anatomical T1 images were acquired using magnetization prepared rapid gradient echo (MPRAGE) sequence: TR=1860ms, TE=2.74ms, FoV=256mm, Voxel size=1×1×1mm, flip angle=8 degrees. Following, functional images were acquired using EPI gradient echo sequence. TR=3000ms, TE=35ms, 44 slices, slice thickness=3 mm, FOV=220mm, Voxel size=2.3×2.3×3mm, flip angle=90 degrees. In total 170 volumes were acquired over the course of the “maternal condition” paradigm. Visual stimuli were displayed to subjects inside the scanner, using a projector (Epson PowerLite 74C, resolution = 1024 × 768), and were back-projected onto a screen mounted above subjects’ heads, and seen by the subjects via an angled mirror. The stimuli were delivered using “Presentation” software (www.neurobs.com).

#### fMRI task

The three maternal conditions paradigm and fMRI sequence began about 50 min after intranasal Oxytocin/Placebo administration. During scanning, participants observed six naturalistic videos of 60 seconds each depicting themselves interacting with their babies (“self” condition) and similar videos of an unfamiliar standard mother interacting with her baby (“other” condition). Between videos a fixation of a black cross over a grey background was presented. Fixation duration was alternated between 15 and 18 seconds. The order of conditions was counterbalanced across subjects. While in the scanner mothers were asked to watch the movies attentively. Video clips were played using VLC media-player (version 2.2 for windows, VideoLAN, France).

#### fMRI Analysis

##### Data preprocessing

Data preprocessing and data analysis were conducted using BrainVoyager QX software package 20.6 (Brain Innovation, Maastricht, The Netherlands) (Goebel et al., 2006). The first 3 functional volumes, before signal stabilization, were automatically discarded by the scanner to allow for T1 equilibrium. Preprocessing of functional scans included 3D motion correction, slice scan time correction, spatial smoothing by a full width at half maximum (FWHM) 6-mm Gaussian kernel, and temporal high-pass filtering. The functional images were then manually aligned and co-registered with 2D anatomical images and incorporated into the 3D datasets through trilinear interpolation. The complete dataset was normalized into Talairach space (TAL) (Talairach and Tournoux, 1988).

##### Whole Brain Analysis

Multi-subject general linear model (GLM) was computed with random effects, with separate subject predictors, in which the various blocks (videos or fixation) were defined as predictors and convoluted with a standard hemodynamic response predictor. Following, a whole brain, three factors (*Maternal Condition* × *Self-Other* × *PBO-OT*) repeated measures ANOVA was performed. Whole brain maps were created and corrected for false discovery rate (FDR) of q<0.050 (Benjamini et al., 1995). For visualization of results, the group contrasts were overlaid on a TAL transformed anatomical brain scan of a single participant.

In order to examine the origin of the significant “maternal condition” factor main effect, we computed group FDR corrected whole brain maps of the contrasts: social≥ unavailable; social≥ unresponsive; unavailable≥ unresponsive. Effects in areas that were not included in our a priori Regions of Interest were reported for descriptive purposes only.

##### Regions-of-Interest Preregistration and Analysis

Region-of-interest (ROI) analysis was conducted on 8 preregistered bilateral defined ROIs (https://osf.io/mszqj/?view_only=0daf10c02c984ead8929452edf44e550) including the amygdala, anterior cingulate (ACC), anterior insula, hippocampus/ parahippocampal gyrus, Temporal pole, VTA, NAcc (all together defined as the “maternal brain”) and the DMN. ROI selection was a priori based on theory and literary meta-reviews (Abraham et al., 2016; Lindquist et al., 2016), and on pilot study of 4 subjects that completed similar paradigm and were not included in the current study. ROIs were defined functionally and anatomically, verified and validated by human brain database platforms: Talairach Daemon (Lancaster et al., 2000) and Neurosynth (Yarkoni et al., 2011), and registered at the Open Science Framework prior to data analysis (OSF, n.d.) (Figure supplement 3).

Beta weights were extracted from ROIs and analyzed with a 3×2×2 (*Maternal Condition* × *Self-Other* × *PBO-OT*) repeated measures ANOVA using SPSS. Thus allowing to investigate main effects of oxytocin administration, and stimulus type and their interactions. In order to further examine the origin of main effects and interactions, simple effect analyses, Scheffé and Bonferroni post hoc tests were conducted. Finally, to investigate brain-behavior coupling, behavioral ratings of mother-infant synchrony during a free-play interaction in the home environment were calculated and entered into Pearson’s correlational analyses together with the extracted values of the ROI’s under Placebo.

##### Within subject correlation (WSC)

In order to test the consistency of temporal patterns during different conditions, between oxytocin and placebo, we calculated a within-subject correlation (WSC) for each subject in each condition in each ROI. The WSC is the Pearson’s correlation for the BOLD time course of each condition (e.g., ‘Self-Social’) in a specific ROI, between the OT and PBO runs. Thus for each participant the correlation indicated how similar was the dynamics of the response in a specific ROI while watching an identical movie clip under the OT or the PBO conditions. Next for each of the ROIs we conducted a 3×2 (*Maternal Condition* × *Self-other*) repeated measures ANOVA. Since no significant self-other main effects, nor self-other × maternal condition interactions were found, we averaged the “self” and “other” variables (containing correlations coefficients) within each condition and ROI. To test differences between the temporal activation consistencies of the preregistered ROIs in the three conditions, a 8×3 (*ROI* × *Maternal Condition*) repeated measures ANOVA was conducted.

#### Behavioral Coding

##### Mirco-level synchrony of the three conditions

To verify that the Social condition indeed was characterized by greater synchrony all video vignettes mothers observed in the scanner were micro-coded by trained coders on a computerized system (Mangold-Interact, Arnstorf, Germany) in 3 seconds frames. Consistent with much prior research in our lab (Feldman and Eidelman, 2007, 2003), four non-verbal categories of infant behavior were coded and each category included a set of mutually exclusive codes (an ‘uncodable’ code was added to each category): Affect (excitement, positive, neutral, medium-fussing, negative, relief after pressure), Gaze (joint attention, to mother’s face, to object or body part, scanning, gaze aversion), Vocalization (no vocalization, positive, neutral, regulatory, negative), and Movement (no movement, hand waving, leg kicking). Mother behavior was coded for Affect and Gaze. Synchrony was defined, consistent with our prior research, by conditional probabilities (infant in state A given mother in state A), indicating episodes when mother and infant were both in social gaze and shared positive affect (Feldman and Eidelman, 2007; Granat et al., 2017).

##### Assessing mother-infant synchrony in the natural ecology

Mother-infant synchrony in the home environment was coded using the Coding Interactive Behavior (CIB) Manual (Feldman, 1998)s. The CIB is a global rating system for adult– child interactions that includes 42 scales that aggregate into theoretically meaningful constructs. The CIB is well-validated with good psychometric properties and has been extensively used across the world in research on health and high-risk population (Feldman, 2012). Consistent with prior research, the synchrony construct of the CIB includes the codes of dyadic reciprocity, mutual adaptation, and fluency, and these codes were averaged into a “mother-infant synchrony” construct. Coding was conducted by a trained coder blind to any other information and inter-rater reliability averaged 95% (k=.87).

#### Statistical analysis

For statistical analysis JASP (Version 0.9 for windows, JASP Team, 2018) and SPSS (SPSS statistics Version 25.0, IBM Corp. Armonk, NY) software were used.

## Acknowledgement

Study was supported by the Simms/Mann Foundation

## References

Abraham E, Hendler T, Shapira-Lichter I, Kanat-Maymon Y, Zagoory-Sharon O, Feldman R. 2014. Father’s brain is sensitive to childcare experiences. Proc Natl Acad Sci U S A 111:9792–7. doi: 10.1073/pnas.1402569111

Abraham E, Hendler T, Zagoory-Sharon O, Feldman R. 2019. Interoception sensitivity in the parental brain during the first months of parenting modulates children’s somatic symptoms six years later: The role of oxytocin. Int J Psychophysiol 136:39–48. doi: 10.1016/j.ijpsycho.2018.02.001

Abraham E, Hendler T, Zagoory-Sharon O, Feldman R. 2016. Network integrity of the parental brain in infancy supports the development of children’s social competencies. Soc Cogn Affect Neurosci 11:1707–1718. doi: 10.1093/scan/nsw090

Abraham E, Raz G, Zagoory-Sharon O, Feldman R. 2018. Empathy networks in the parental brain and their long-term effects on children’s stress reactivity and behavior adaptation. Neuropsychologia 116:75–85. doi: 10.1016/j.neuropsychologia.2017.04.015

Allen M, Tsakiris M. 2018. The body as first prior : Interoceptive predictive processing and the primacy of self-models In: Tsakiris M, De Preester H, editors. The Interoceptive Mind From Homeostasis to Awareness. Oxford University Press. pp. 27–45.

Anders S, Heinzle J, Weiskopf N, Ethofer T, Haynes JD. 2011. Flow of affective information between communicating brains. Neuroimage 54:439–446. doi: 10.1016/j.neuroimage.2010.07.004

Andrews-Hanna JR, Smallwood J, Spreng RN. 2014. The default network and self-generated thought: Component processes, dynamic control, and clinical relevance. Ann N Y Acad Sci 1316:29–52. doi: 10.1111/nyas.12360

Araujo HF, Kaplan J, Damasio H, Damasio A. 2015. Neural correlates of different self domains. Brain Behav 5:n/a-n/a. doi: 10.1002/brb3.409

Atzil S, Gao W, Fradkin I, Barrett LF. 2018. Growing a social brain. Nat Hum Behav 2:624–636. doi: 10.1038/s41562-018-0384-6

Atzil S, Hendler T, Feldman R. 2014. The brain basis of social synchrony. Soc Cogn Affect Neurosci 9:1193–202. doi: 10.1093/scan/nst105

Atzil S, Hendler T, Feldman R. 2011. Specifying the neurobiological basis of human attachment: brain, hormones, and behavior in synchronous and intrusive mothers. Neuropsychopharmacology 36:2603–15. doi: 10.1038/npp.2011.172

Báez-Mendoza R, Schultz W. 2013. The role of the striatum in social behavior. Front Neurosci 7:233. doi: 10.3389/fnins.2013.00233

Bartz JA, Zaki J, Bolger N, Ochsner KN. 2011. Social effects of oxytocin in humans: Context and person matter. Trends Cogn Sci 15:301–309. doi: 10.1016/j.tics.2011.05.002

Benjamini Y, Hochberg Y, Benjamini, Yoav HY. 1995. Benjamini and Y FDR.pdf. J R Stat Soc Ser B. doi: 10.2307/2346101

Carter CS. 2014. Oxytocin Pathways and the Evolution of Human Behavior. Annu Rev Psychol 65:17–39. doi: 10.1146/annurev-psych-010213-115110

Champagne F, Diorio J, Sharma S, Meaney MJ. 2001. Naturally occurring variations in maternal behavior in the rat are associated with differences in estrogen-inducible central oxytocin receptors. Proc Natl Acad Sci U S A. doi: 10.1073/pnas.221224598

Chen X, Gautam P, Haroon E, Rilling JK. 2017. Within vs. between-subject effects of intranasal oxytocin on the neural response to cooperative and non-cooperative social interactions. Psychoneuroendocrinology 78:22–30. doi: 10.1016/j.psyneuen.2017.01.006

Cohn JF, Tronick EZ. 1988. Mother-Infant Face-to-Face Interaction: Influence is Bidirectional and Unrelated to Periodic Cycles in Either Partner’s Behavior. Dev Psychol 24:386–392. doi: 10.1037/0012-1649.24.3.386

Cox JL, Holden JM, Sagovsky R. 1987. Detection of postnatal depression. Development of the 10-item Edinburgh Postnatal Depression Scale. Br J Psychiatry 150:782–786. doi: 10.1192/bjp.150.6.782

Craig AD. 2009. How do you feel - now? The anterior insula and human awareness. Nat Rev Neurosci. doi: 10.1038/nrn2555

Davey CG, Pujol J, Harrison BJ. 2016. Mapping the self in the brain’s default mode network. Neuroimage 132:390–397. doi: 10.1016/j.neuroimage.2016.02.022

Davis EP, Stout SA, Molet J, Vegetabile B, Glynn LM, Sandman CA, Heins K, Stern H, Baram TZ. 2017. Exposure to unpredictable maternal sensory signals influences cognitive development across species. Proc Natl Acad Sci U S A 114:10390–10395. doi: 10.1073/pnas.1703444114

Elmadih A, Wan MW, Downey D, Elliott R, Swain JE, Abel KM. 2016. Natural variation in maternal sensitivity Is reflected in maternal brain responses to infant stimuli. Behav Neurosci 130:500–510. doi: 10.1037/bne0000161

Feldman R. 2020. What is resilience: an affiliative neuroscience approach. World Psychiatry 19:132–150. doi: 10.1002/wps.20729

Feldman R. 2017. The Neurobiology of Human Attachments. Trends Cogn Sci 21:80–99. doi: 10.1016/j.tics.2016.11.007

Feldman R. 2016. The neurobiology of mammalian parenting and the biosocial context of human caregiving. Horm Behav 77:3–17. doi: 10.1016/j.yhbeh.2015.10.001

Feldman R. 2015a. The adaptive human parental brain: implications for children’s social development. Trends Neurosci 38:387–399. doi: 10.1016/J.TINS.2015.04.004

Feldman R. 2015b. Sensitive periods in human social development: New insights from research on oxytocin, synchrony, and high-risk parenting. Dev Psychopathol 27:369–395. doi: 10.1017/S0954579415000048

Feldman R. 2012. Parenting Behavior as the Environment Where Children Grow, The Cambridge Handbook of Environment in Human Development. doi: 10.1017/cbo9781139016827.031

Feldman R. 2010. The relational basis of adolescent adjustment: trajectories of mother–child interactive behaviors from infancy to adolescence shape adolescents’ adaptation. Attach Hum Dev 12:173–192. doi: 10.1080/14616730903282472

Feldman R. 2007. Parent-infant synchrony and the construction of shared timing; physiological precursors, developmental outcomes, and risk conditions. J Child Psychol Psychiatry Allied Discip 48:329–354. doi: 10.1111/j.1469-7610.2006.01701.x

Feldman R. 1998. Coding Interactive Behavior Manual. Tel Aviv, Israel: Bar-Ilan Univ Press.

Feldman R, Eidelman AI. 2007. Maternal Postpartum Behavior and the Emergence of Infant–Mother and Infant–Father Synchrony in Preterm and Full-Term Infants: The Role of Neonatal Vagal Tone. Dev Psychobiol 49:290–302.

Feldman R, Eidelman AI. 2003. Direct and indirect effects of breast milk on the neurobehavioral and cognitive development of premature infants. Dev Psychobiol 43:109–119. doi: 10.1002/dev.10126

Feldman R, Gordon I, Zagoory-Sharon O. 2011a. Maternal and paternal plasma, salivary, and urinary oxytocin and parent-infant synchrony: Considering stress and affiliation components of human bonding. Dev Sci 14:752–761. doi: 10.1111/j.1467-7687.2010.01021.x

Feldman R, Gordon I, Zagoory-Sharon O. 2010. The cross-generation transmission of oxytocin in humans. Horm Behav 58:669–676. doi: 10.1016/j.yhbeh.2010.06.005

Feldman R, Magori-Cohen R, Galili G, Singer M, Louzoun Y. 2011b. Mother and infant coordinate heart rhythms through episodes of interaction synchrony. Infant Behav Dev 34:569–577. doi: 10.1016/j.infbeh.2011.06.008

Fotopoulou A, Tsakiris M. 2017. Mentalizing homeostasis: The social origins of interoceptive inference 4145. doi: 10.1080/15294145.2017.1294031

Francis D, Diorio J, Liu D, Meaney MJ. 1999. Nongenomic transmission across generations of maternal behavior and stress responses in the rat. Science (80-) 286:1155–1158. doi: 10.1126/science.286.5442.1155

Goebel R, Esposito F, Formisano E. 2006. Analysis of Functional Image Analysis Contest (FIAC) data with BrainVoyager QX: From single-subject to cortically aligned group General Linear Model analysis and self-organizing group Independent Component Analysis. Hum Brain Mapp 27:392–401. doi: 10.1002/hbm.20249

González-Mariscal G. 2007. Mother rabbits and their offspring: Timing is everything. Dev Psychobiol. doi: 10.1002/dev.20196

Gordon I, Vander Wyk BC, Bennett RH, Cordeaux C, Lucas M V., Eilbott JA, Zagoory-Sharon O, Leckman JF, Feldman R, Pelphrey KA. 2013. Oxytocin enhances brain function in children with autism. Proc Natl Acad Sci U S A 110:20953–20958. doi: 10.1073/pnas.1312857110

Grace SA, Rossell SL, Heinrichs M, Kordsachia C, Labuschagne I. 2018. Oxytocin and brain activity in humans: A systematic review and coordinate-based meta-analysis of functional MRI studies. Psychoneuroendocrinology 96:6–24. doi: 10.1016/j.psyneuen.2018.05.031

Granat A, Gadassi R, Gilboa-Schechtman E, Feldman R. 2017. Maternal depression and anxiety, social synchrony, and infant regulation of negative and positive emotions. Emotion 17:11–27. doi: 10.1037/emo0000204

Gray SAO, Jones CW, Theall KP, Glackin E, Drury SS. 2017. Thinking Across Generations: Unique Contributions of Maternal Early Life and Prenatal Stress to Infant Physiology. J Am Acad Child Adolesc Psychiatry 56:922–929. doi: 10.1016/j.jaac.2017.09.001

Gur R, Tendler A, Wagner S. 2014. Long-Term Social Recognition Memory Is Mediated by Oxytocin-Dependent Synaptic Plasticity in the Medial Amygdala. Biol Psychiatry 76:377–386. doi: 10.1016/J.BIOPSYCH.2014.03.022

Halevi G, Djalovski A, Kanat-Maymon Y, Yirmiya K, Zagoory-Sharon O, Koren L, Feldman R. 2017. The social transmission of risk: Maternal stress physiology, synchronous parenting, and well-being mediate the effects of war exposure on child psychopathology. J Abnorm Psychol 126:1087–1103. doi: 10.1037/abn0000307

Hammock EAD. 2015. Developmental perspectives on oxytocin and vasopressin. Neuropsychopharmacology 40:24–42. doi: 10.1038/npp.2014.120

Hasson U, Nir Y, Levy I, Fuhrmann G, Malach R. 2004. Intersubject synchronization of cortical activity during natural vision. Science (80-) 303:1634–1640.

Hayes LD. 2000. To nest communally or not to nest communally: A review of rodent communal nesting and nursing. Anim Behav 59:677–688. doi: 10.1006/anbe.1999.1390

Higashida H, Lopatina O, Yoshihara T, Pichugina YA, Soumarokov AA, Munesue T, Minabe Y, Kikuchi M, Ono Y, Korshunova N, Salmina AB. 2010. Oxytocin signal and social behaviour: Comparison among adult and infant oxytocin, oxytocin receptor and CD38 gene knockout mice. J Neuroendocrinol 22:373–379. doi: 10.1111/j.1365-2826.2010.01976.x

Hofer M a. 1994. Hidden Regulators in Attachment, Separation, and Loss Myron A. Hofer Monographs of the Society for Research in Child Development, Vol. 59, No. 2 / 3, The Development of Emotion Regulation : Biological and Behavioral Considerations. (1994), pp. Emotion 59:192–207.

Hofer MA. 1987. Early Social Relationships : A Psychobiologist’ s View Author (s): Myron A. Hofer Published by : Wiley on behalf of the Society for Research in Child Development Stable URL : http://www.jstor.org/stable/1130203 REFERENCES Linked references are availab. Child Dev 58:633–647.

Insel TR, Young LJ. 2001. The neurobiology of attachment. Nat Rev Neurosci 2:129–136. doi: 10.1038/35053579

Kim P, Capistrano CG, Erhart A, Gray-Schiff R, Xu N. 2017. Socioeconomic disadvantage, neural responses to infant emotions, and emotional availability among first-time new mothers. Behav Brain Res 325:188–196. doi: 10.1016/j.bbr.2017.02.001

Kim P, Rigo P, Leckman JF, Mayes LC, Cole PM, Feldman R, Swain JE. 2015. A prospective longitudinal study of perceived infant outcomes at 18-24 months: Neural and psychological correlates of parental thoughts and actions assessed during the first month postpartum. Front Psychol 6:1–14. doi: 10.3389/fpsyg.2015.01772

Kim P, Strathearn L, Swain JE. 2016. The maternal brain and its plasticity in humans. Horm Behav 77:113–123. doi: 10.1016/j.yhbeh.2015.08.001

Kinreich S, Djalovski A, Kraus L, Louzoun Y, Feldman R. 2017. Brain-to-Brain Synchrony during Naturalistic Social Interactions. Sci Rep 7:1–12. doi: 10.1038/s41598-017-17339-5

Kringelbach ML, Lehtonen A, Squire S, Harvey AG, Craske MG, Holliday IE, Green AL, Aziz TZ, Hansen PC, Cornelissen PL, Stein A. 2008. A specific and rapid neural signature for parental instinct. PLoS One 3. doi: 10.1371/journal.pone.0001664

Krol KM, Moulder RG, Lillard TS, Grossmann T, Connelly JJ. 2019. Epigenetic dynamics in infancy and the impact of maternal engagement. Sci Adv 5:1–8. doi: 10.1126/sciadv.aay0680

Kundakovic M, Champagne FA. 2015. Early-life experience, Epigenetics, and the developing brain. Neuropsychopharmacology 40:141–153. doi: 10.1038/npp.2014.140

Lancaster JL, Woldorff MG, Parsons LM, Liotti M, Freitas CS, Rainey L, Kochunov P V., Nickerson D, Mikiten SA, Fox PT. 2000. Automated Talairach Atlas labels for functional brain mapping. Hum Brain Mapp 10:120–131. doi: 10.1002/1097-0193(200007)10:3<120::AID-HBM30>3.0.CO;2-8

Leong V, Byrne E, Clackson K, Georgieva S, Lam S, Wass S. 2017. Speaker gaze increases information coupling between infant and adult brains. Proc Natl Acad Sci U S A 114:13290–13295. doi: 10.1073/pnas.1702493114

Levy J, Goldstein A, Feldman R. 2019. The neural development of empathy is sensitive to caregiving and early trauma. Nat Commun 10. doi: 10.1038/s41467-019-09927-y

Lindquist KA, Satpute AB, Wager TD, Weber J, Barrett LF. 2016. The Brain Basis of Positive and Negative Affect: Evidence from a Meta-Analysis of the Human Neuroimaging Literature. Cereb Cortex 26:1910–1922. doi: 10.1093/cercor/bhv001

Lopatina O, Inzhutova A, Salmina AB, Higashida H. 2012. The roles of oxytocin and CD38 in social or parental behaviors. Front Neurosci 6:1–12. doi: 10.3389/fnins.2012.00182

Lucion AB, Bortolini MC. 2014. Mother-pup interactions: Rodents and humans. Front Endocrinol (Lausanne) 5:1–5. doi: 10.3389/fendo.2014.00017

Marlin BJ, Mitre M, D’Amour JA, Chao M V, Froemke RC. 2015. Oxytocin enables maternal behaviour by balancing cortical inhibition. Nature 520:499–504. doi: 10.1038/nature14402

Martins DA, Mazibuko N, Zelaya F, Vasilakopoulou S, Loveridge J, Oates A, Maltezos S, Mehta M, Wastling S, Howard M, McAlonan G, Murphy D, Williams SCR, Fotopoulou A, Schuschnig U, Paloyelis Y. 2020. Effects of route of administration on oxytocin-induced changes in regional cerebral blood flow in humans. Nat Commun 11:1–16. doi: 10.1038/s41467-020-14845-5

Meaney MJ. 2001. MATERNAL CARE,GENE EXPRESSION, AND THE TRANSMISSION OF INDIVIDUAL DIFFERENCES IN STRESS REACTIVITY ACROSS GENERATIONS. Annu Rev Neurosci 24:1161–1192. doi: 10.1146/annurev.neuro.24.1.1161

Naber F, van IJzendoorn MH, Deschamps P, van Engeland H, Bakermans-Kranenburg MJ. 2010. Intranasal oxytocin increases fathers’ observed responsiveness during play with their children: A double-blind within-subject experiment. Psychoneuroendocrinology 35:1583–1586. doi: 10.1016/j.psyneuen.2010.04.007

Neumann ID. 2008. Brain Oxytocin: A Key Regulator of Emotional and Social Behaviours in Both Females and Males. J Neuroendocrinol 20:858–865. doi: 10.1111/j.1365-2826.2008.01726.x

Neumann ID, Slattery DA. 2016. Oxytocin in General Anxiety and Social Fear: A Translational Approach. Biol Psychiatry 79:213–221. doi: 10.1016/j.biopsych.2015.06.004

Noriuchi M, Kikuchi Y, Senoo A. 2008. The Functional Neuroanatomy of Maternal Love: Mother’s Response to Infant’s Attachment Behaviors. Biol Psychiatry 63:415–423. doi: 10.1016/j.biopsych.2007.05.018

Northoff G, Heinzel A, de Greck M, Bermpohl F, Dobrowolny H, Panksepp J. 2006. Self-referential processing in our brain-A meta-analysis of imaging studies on the self. Neuroimage 31:440–457. doi: 10.1016/j.neuroimage.2005.12.002

Numan M. 2020. The Parental Brain: Mechanisms, Development, and Evolutiontle. Oxford University Press.

Numan M, Young LJ. 2016. Neural mechanisms of mother–infant bonding and pair bonding: Similarities, differences, and broader implications. Horm Behav 77:98–112. doi: 10.1016/j.yhbeh.2015.05.015

Nummenmaa L, Glerean E, Viinikainen M, Jääskeläinen IP, Hari R, Sams M. 2012. Emotions promote social interaction by synchronizing brain activity across individuals. Proc Natl Acad Sci U S A 109:9599–9604. doi: 10.1073/pnas.1206095109

Oettl LL, Ravi N, Schneider M, Scheller MF, Schneider P, Mitre M, da Silva Gouveia M, Froemke RC, Chao M V., Young WS, Meyer-Lindenberg A, Grinevich V, Shusterman R, Kelsch W. 2016. Oxytocin Enhances Social Recognition by Modulating Cortical Control of Early Olfactory Processing. Neuron 90:609–621. doi: 10.1016/j.neuron.2016.03.033

Olazábal DE, Young LJ. 2006. Oxytocin receptors in the nucleus accumbens facilitate “spontaneous” maternal behavior in adult female prairie voles. Neuroscience 141:559–568. doi: 10.1016/j.neuroscience.2006.04.017

OSF. n.d. OSF, https://osf.io/myprojects/. https://osf.io/myprojects/

Park HD, Bernasconi F, Bello-Ruiz J, Pfeiffer C, Salomon R, Blanke O. 2016. Transient modulations of neural responses to heartbeats covary with bodily self-consciousness. J Neurosci 36:8453–8460. doi: 10.1523/JNEUROSCI.0311-16.2016

Peer M, Salomon R, Goldberg I, Blanke O, Arzy S. 2015. Brain system for mental orientation in space, time, and person. Proc Natl Acad Sci U S A 112:11072–11077. doi: 10.1073/pnas.1504242112

Pratt M, Goldstein A, Feldman R. 2018. Child brain exhibits a multi-rhythmic response to attachment cues. Soc Cogn Affect Neurosci 13:957–966. doi: 10.1093/scan/nsy062

Pratt M, Zeev-Wolf M, Goldstein A, Feldman R. 2019. Exposure to early and persistent maternal depression impairs the neural basis of attachment in preadolescence. Prog Neuro-Psychopharmacology Biol Psychiatry 93:21–30. doi: 10.1016/j.pnpbp.2019.03.005

Rassovsky Y, Harwood A, Zagoory-Sharon O, Feldman R. 2019. Martial arts increase oxytocin production. Sci Rep 9:12980. doi: 10.1038/s41598-019-49620-0

Rilling JK, Mascaro JS. 2017. The neurobiology of fatherhood. Curr Opin Psychol 15:26–32. doi: 10.1016/j.copsyc.2017.02.013

Ross HE, Cole CD, Smith Y, Neumann ID, Landgraf R, Murphy AZ, Young LJ. 2009. Characterization of the oxytocin system regulating affiliative behavior in female prairie voles. Neuroscience 162:892–903. doi: 10.1016/j.neuroscience.2009.05.055

Ross HE, Young LJ. 2009. Oxytocin and the neural mechanisms regulating social cognition and affiliative behavior. Front Neuroendocrinol 30:534–547. doi: 10.1016/j.yfrne.2009.05.004

Russell AF, Brotherton PNM, McIlrath GM, Sharpe LL, Clutton-Brock TH. 2003. Breeding success in cooperative meerkats: Effects of helper number and maternal state. Behav Ecol 14:486–492. doi: 10.1093/beheco/arg022

Salomon R, Levy DR, Malach R. 2014. Deconstructing the default: Cortical subdivision of the default mode/intrinsic system during self-related processing. Hum Brain Mapp 35:1491–1502. doi: 10.1002/hbm.22268

Salomon R, Malach R, Lamy D. 2009. Involvement of the intrinsic/default system in movement-related self recognition. PLoS One 4. doi: 10.1371/journal.pone.0007527

Salomon R, Ronchi R, Dönz J, Bello-Ruiz J, Herbelin B, Martet R, Faivre N, Schaller K, Blanke O. 2016. The insula mediates access to awareness of visual stimuli presented synchronously to the heartbeat. J Neurosci 36:5115–5127. doi: 10.1523/JNEUROSCI.4262-15.2016

Schirmer A, Meck WH, Penney TB. 2016. The Socio-Temporal Brain: Connecting People in Time. Trends Cogn Sci 20:760–772. doi: 10.1016/j.tics.2016.08.002

Schulkin J, Sterling P. 2019. Allostasis: A Brain-Centered, Predictive Mode of Physiological Regulation. Trends Neurosci 42:740–752. doi: 10.1016/j.tins.2019.07.010

Schultz W. 2016. Reward functions of the basal ganglia. J Neural Transm 123:679–693.

Seth AK. 2013. Interoceptive inference, emotion, and the embodied self. Trends Cogn Sci. doi: 10.1016/j.tics.2013.09.007

Seth AK, Friston KJ. 2016. Active interoceptive inference and the emotional brain. Philos Trans R Soc B Biol Sci 371. doi: 10.1098/rstb.2016.0007

Shamay-Tsoory SG, Abu-Akel A. 2016. The Social Salience Hypothesis of Oxytocin. Biol Psychiatry 79:194–202. doi: 10.1016/j.biopsych.2015.07.020

Sokolowski K, Corbin JG. 2012. Wired for behaviors: from development to function of innate limbic system circuitry. Front Mol Neurosci 5:55. doi: 10.3389/fnmol.2012.00055

Sterling P. 2012. Allostasis: A model of predictive regulation. Physiol Behav 106:5–15. doi: 10.1016/j.physbeh.2011.06.004

Swain JE, Kim P, Spicer J, Ho SS, Dayton CJ, Elmadih A, Abel KM. 2014. Approaching the biology of human parental attachment: Brain imaging, oxytocin and coordinated assessments of mothers and fathers. Brain Res 1580:78–101. doi: 10.1016/j.brainres.2014.03.007

Talairach J, Tournoux P. 1988. A Coplanar Stereotaxic Atlas of a Human Brain. Thieme, Stuttgart.

Tronick EZ. 1989. Emotions and emotional communication in infants. Am Psychol 44:112–119. doi: 10.1037//0003-066x.44.2.112

Ulmer-Yaniv A, Salomon R, Waidergoren S, Shimon-Raz O, Djalovski A, Feldman R. 2020. Synchronous Caregiving from Birth to Adulthood Tunes Humans’ Social Brain Authors:

Valtcheva S, Froemke RC. 2019. Neuromodulation of maternal circuits by oxytocin. Cell Tissue Res 375:57–68. doi: 10.1007/s00441-018-2883-1

Walum H, Young LJ. 2018. The neural mechanisms and circuitry of the pair bond. Nat Rev Neurosci 19:643–654. doi: 10.1038/s41583-018-0072-6

Wang D, Yan X, Li M, Ma Y. 2017. Neural substrates underlying the effects of oxytocin: A quantitative meta-analysis of pharmaco-imaging studies. Soc Cogn Affect Neurosci 12:1565–1573. doi: 10.1093/scan/nsx085

Weisman O, Zagoory-Sharon O, Feldman R. 2012a. Oxytocin administration to parent enhances infant physiological and behavioral readiness for social engagement. Biol Psychiatry 72:982–989. doi: 10.1016/j.biopsych.2012.06.011

Weisman O, Zagoory-Sharon O, Feldman R. 2012b. Intranasal oxytocin administration is reflected in human saliva. Psychoneuroendocrinology 37:1582–1586. doi: 10.1016/j.psyneuen.2012.02.014

Wigton R, Radua J, Allen P, Averbeck B, Meyer-Lindenberg A, McGuire P, Sukhi S, Fusar-Poli P. 2015. Neurophysiological effects of acute oxytocin administration: Systematic review and meta-analysis of placebo-controlled imaging studies. J Psychiatry Neurosci 40:E1–E22. doi: 10.1503/jpn.130289

Wittmann M. 2013. The inner sense of time: How the brain creates a representation of duration. Nat Rev Neurosci 14:217–223. doi: 10.1038/nrn3452

Yarkoni T, Poldrack RA, Nichols TE, Van Essen DC, Wager TD. 2011. Large-scale automated synthesis of human functional neuroimaging data. Nat Methods 8:665–670. doi: 10.1038/nmeth.1635

Zink CF, Meyer-Lindenberg A. 2012. Human neuroimaging of oxytocin and vasopressin in social cognition. Horm Behav 61:400–409. doi: 10.1016/j.yhbeh.2012.01.016

