## Supplementary Materials for "Mother Brain is Wired for Social Moments"

**This file includes:**

Figures supplement 1 to supplement 3

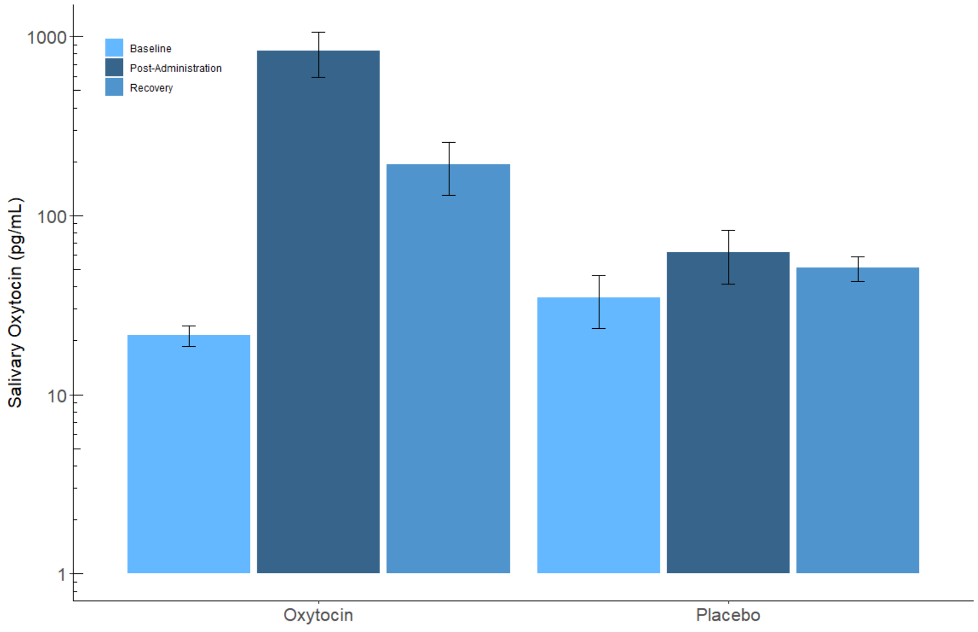


**Figure supplement 1.** Mother salivary oxytocin levels (pg/mL) in the oxytocin and placebo conditions. 2×3 Repeated measures ANOVA revealed a significant *PBO-OT* × *Time* interaction effect, *PBO-OT* main effect and *Time* main effect, showing marked increase in OT level following OT administration. In the placebo condition, no such increase was observed. All effects were Greenhouse-Geisser corrected. Error bars represent standard error of the mean.

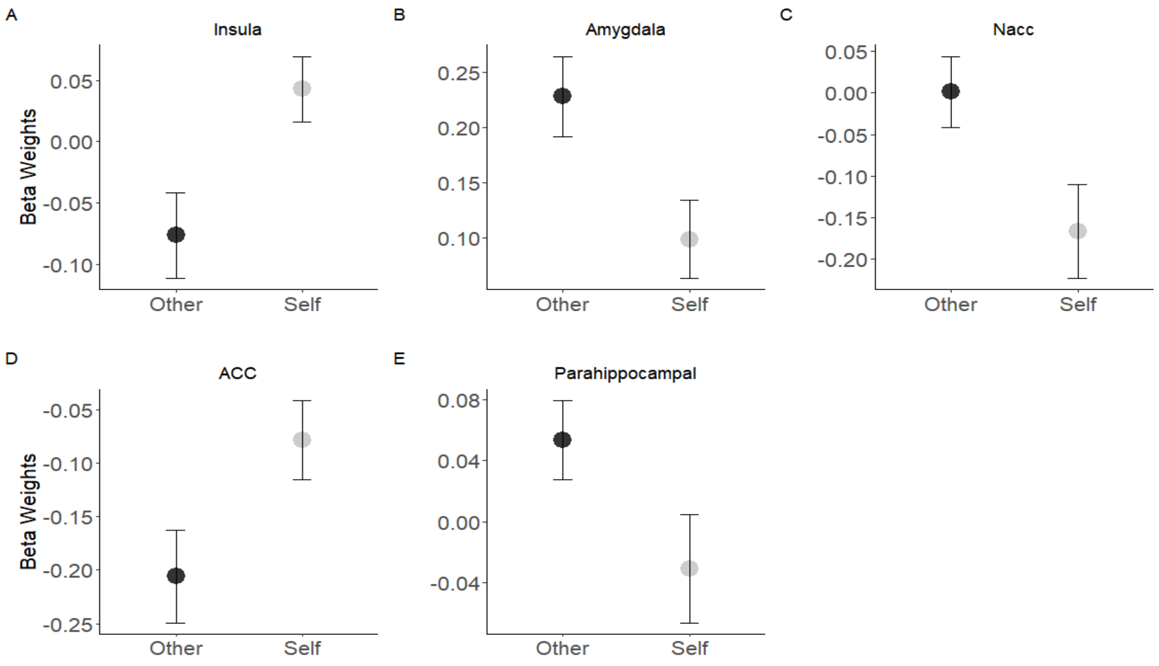


**Figure supplement 2.** *Self-Other* main effect in five ROIs of the maternal brain. 3 factorial repeated measures ANOVA (*Maternal Condition*× *Self-Other*× *PBO-OT*) was performed on the beta values extracted from each of the ROIs. The insula and ACC showed greater activation in response to *self*-stimuli (grey color) while greater activation in response to *other*- stimuli was found in the amygdala, parahippocampal gyrus and in the NAcc. Bars depict Standard error of the mean. NAcc, Nucleus accumbens; parahippocampal, parahippocampal gyrus.

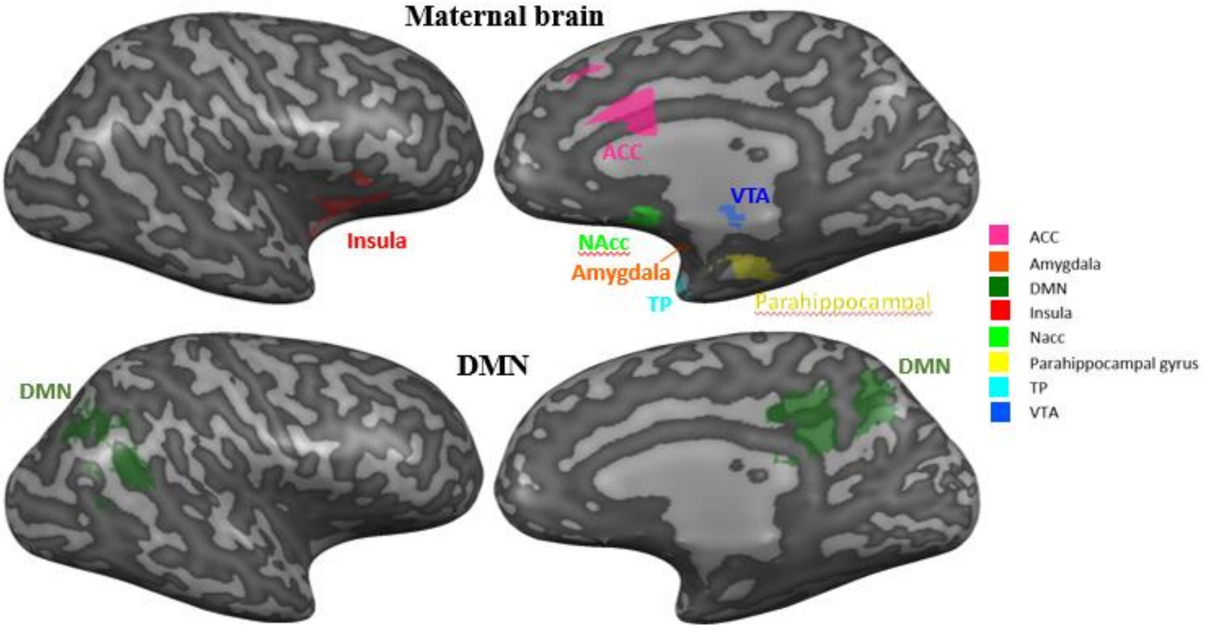


**Figure supplement 3**. The "Maternal Brain" and the DMN. 8 preregistered ROI's laid on right hemisphere. ACC, Anterior cingulate; DMN, Default mode network; NAcc, Nucleus accumbens; TP, Temporal pole; VTA, ventral tegmental area.
